# moPPIt: *De Novo* Generation of Motif-Specific and Functionally Active Peptide Binders via Discrete Flow Matching

**DOI:** 10.1101/2024.07.31.606098

**Authors:** Tong Chen, Zachary Quinn, Kunal Mishra, Erin C. O’Connor, Sophia E. Silver, Yinuo Zhang, Mary Jo Valencia, Ying Mei, Jacques Behmoaras, Leonardo M.R. Ferreira, Pranam Chatterjee

## Abstract

Targeting specific functional motifs, whether conserved viral epitopes, intrinsically disordered regions (IDRs), or fusion breakpoints, is essential for modulating protein function and protein-protein interactions (PPIs). Current design methods, however, depend on stable tertiary structures, limiting their utility for disordered or dynamic targets. Here, we present a **mo**tif-specific **PPI t**argeting algorithm (**moPPIt)**, a framework for the *de novo* generation of motif-specific peptide binders derived solely from target sequence data. The core of this approach is BindEvaluator, a transformer architecture that interpolates protein language model embeddings to predict peptide-protein binding site interactions with high accuracy (AUC = 0.97). We integrate this predictor into a novel **M**ulti-**O**bjective-**G**uided **D**iscrete **F**low **M**atching (**MOG-DFM**) framework, which steers generative trajectories toward peptides that simultaneously maximize binding affinity and motif specificity. After comprehensive *in silico* validation of binding and motif-specific targeting, we validate moPPIt *in vitro* by generating binders that strictly discriminate between the FN3 and IgG domains of NCAM1, confirming domain-level specificity, and further demonstrate precise targeting of IDRs by generating binders to the N-terminal disordered domain of β-catenin. In functional, disease-relevant assays, moPPIt-designed peptides specifically targeting the GM-CSF receptor ɑ subunit effectively block human macrophage polarization. Finally, we demonstrate utility in cell engineering, where binders directed against a defined motif on a synthetic cell surface ligand (AGR2t) drive specific chimeric antigen receptor regulatory T cell (CAR Treg) activation and suppressive function, including in the context of human induced pluripotent stem cell (hiPSC)-derived cardiomyocytes. Altogether, moPPIt serves as a theoretically-justified, sequence-based paradigm for controllably targeting the complete proteome with immediate therapeutic applications.

## Introduction

Motif-specific targeting of protein-protein interactions (PPIs) offers the potential for highly selective biotherapeutics that can modulate protein function while minimizing off-target effects.^1^ This precision is particularly critical for “undruggable” targets where specificity is dictated by transient or localized features rather than deep hydrophobic pockets. These include the breakpoint junctions of fusion oncoproteins like PAX3::FOXO1 and EWS::FLI1,^2^ the allosteric regulatory sites of membrane proteins such as GPCRs,^3,4^ and the aggregation-prone regions of proteins like tau, α-synuclein, and GFAP involved in neurodegenerative diseases.^5,6^ In these contexts, the therapeutic goal is not merely to bind the protein, but to selectively engage specific functional epitopes to modulate a distinct biological state.

While computational design has accelerated binder discovery, current state-of-the-art methods like RFDiffusion, BindCraft, and BoltzGen operate fundamentally in structure space.^7–9^ This reliance on static coordinate representations creates a significant bottleneck for targets lacking stable tertiary conformations, such as intrinsically disordered proteins (IDPs) or flexible loops, which are absent from structural training sets.^10,11^ More critically, structure-based models often focus on local geometric complementarity at the expense of global protein properties (such as solubility, stability, immunogenicity, global specificity, and synthesis feasibility).^12^ Consequently, there remains a critical need for generative frameworks that can design binders *de novo* without requiring a fixed target backbone, while simultaneously optimizing for multiple biophysical objectives.

We have previously developed protein language models (pLMs), including SaLT&PepPr, PepPrCLIP, and PepMLM, to generate therapeutic peptides directly from sequence information.^13–15^ While these models successfully produce binders, including those validated *in vivo*,^16^ they lack the ability to explicitly incorporate granular design constraints, such as motif specificity or binding affinity, at inference time. To enable this controllability, we have recently developed guidance frameworks on discrete diffusion models, including PepTune, TR2-D2, and DDPP.^17–19^ More recently, we have extended this to discrete flow matching, such with base models including PepDFM and Gumbel-Softmax Flow Matching^20,21^ and guidance frameworks, MOG-DFM and AReUReDi.^20,22^ All of these strategies allow for the precise, property-guided steering of the generative process of peptides. However, we have yet to couple controllable generation of peptides with the specific objective of motif targeting, leaving us unable to direct binders to distinct functional epitopes on a target surface.

To address this, we present a <u>mo</u>tif-specific <u>PPI t</u>argeting algorithm, termed **moPPIt**, which bridges this gap by enabling the *de novo* design of optimized, motif-specific binders derived solely from target sequence data. moPPIt extends our Multi-Objective-Guided Discrete Flow Matching (MOG-DFM) framework^20^ to steer the generative trajectory of PepDFM using two complementary guidance signals: a pre-trained affinity predictor to ensure binding strength, and BindEvaluator, a novel transformer architecture we have developed to enforce motif specificity. By interpolating pLM embeddings, BindEvaluator accurately predicts the binding sites of specific peptide-protein interactions by capturing both local residue-level interactions and global sequence dependencies, allowing moPPIt to direct generation toward specific epitopes without structural priors.

We validate moPPIt through a series of increasingly complex biological challenges that highlight the versatility of this sequence-based design framework. We first demonstrate domain-level specificity *in vitro* by designing binders that strictly discriminate between the fibronectin type III (FN3) and immunoglobulin domains of NCAM1.^23^ Extending this to the disordered proteome, we generate and validate binders to the N-terminal intrinsically disordered region (IDR) of β-catenin.^24,25^ Finally, we demonstrate therapeutic utility in functional cell engineering contexts: moPPIt-designed peptides specifically targeting the GM-CSF receptor ɑ subunit, potently block macrophage polarization,^26,27^ and binders directed against a synthetic cell surface ligand (AGR2t) drive specific CAR T-regulatory cell activation and suppressive function, including in the context of human induced pluripotent stem cell-derived cardiomyocytes.^28^ Collectively, these results establish moPPIt as a comprehensive, purely sequence-based paradigm for controllably targeting the complete proteome.

## Results

### BindEvaluator accurately predicts target binding sites provided two interacting sequences

To enable motif-specific generation, we first developed BindEvaluator to predict peptide-protein binding sites from sequence inputs (**Figure 1A**). Binder and target sequences are embedded via the pretrained ESM-2-650M pLM,^29^ after which target embeddings are processed by a dilated convolutional neural network (CNN) to capture local residue features. These embeddings then pass through multi-head attention layers to encode global dependencies, while reciprocal attention modules integrate target and binder representations to explicitly model their interaction. Finally, the fused representation is processed by feed-forward layers to predict the specific binding residues.

**Figure 1:**
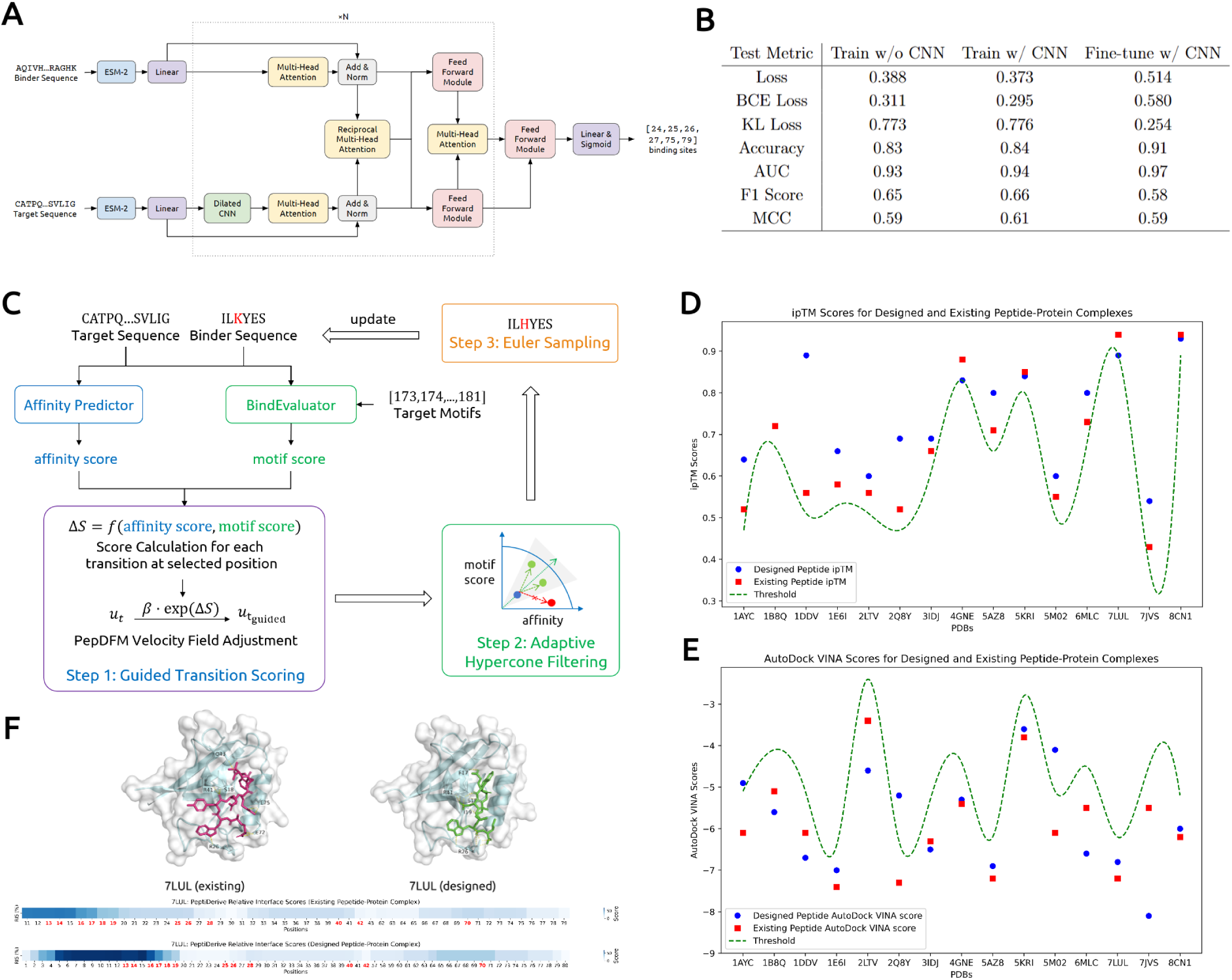
Overview of BindEvaluator and moPPIt. **(A) The architecture of BindEvaluator.** BindEvaluator predicts the binding residues on the target protein given a target sequence and a binder sequence. The binder and target sequences are first processed using a pre-trained ESM-2-650M model to obtain their embeddings. The target sequence embeddings are further refined using a dilated CNN module to capture local features. Both embeddings are then passed through multi-head attention modules to capture global dependencies. Reciprocal multi-head attention modules integrate the representations of the target and binder sequences, allowing for the capture of binder-target interaction information. Feed-forward and linear layers subsequently process the refined embeddings to predict the binding sites. **(B) Test performance metrics of BindEvaluator across different training configurations.** Performance metrics were calculated for BindEvaluator across three configurations: trained without dilated CNN modules, trained with dilated CNN modules, and fine-tuned for peptide-protein binding site prediction. Metrics include overall loss, binary cross-entropy (BCE) loss, KL divergence loss, accuracy, area under the ROC curve (AUC), F1 score, and Matthews correlation coefficient (MCC). **(C) Schematic of moPPIt.** The algorithm starts with a randomly initialized binder sequence as well as a target protein sequence and target motifs. MOG-DFM framework is used to iteratively update the binder sequence so as to optimize motif specificity and binding affinity in the state space. **(D), (E) Hit rate of moPPIt on structured targets with known binders for ipTM and AutoDock VINA scores.** The scores for known peptides (red) from PDB structures were compared to moPPIt-designed peptides (blue) for the same target proteins. An ipTM score below 0.05 of the existing peptide (green line) was used as a threshold to call hits. An AutoDock VINA score above 1.0 of the existing peptide (green line) was used as the threshold to call hits. **(F)** AutoDock VINA docking visualization of protein (PDB ID: 7LUL) with existing and designed peptide binders, highlighting interacting residues.

We initially trained a baseline model without dilated CNNs on over 500,000 annotated protein-protein interactions (PPIs) from the PPIRef dataset.^30^ While this model effectively distinguished binding residues, we hypothesized that dilated CNNs would enhance the extraction of local binding features. Indeed, incorporating CNN modules yielded observable improvements across test metrics (**Figure 1B, Supplementary Figures 1A-B**). To adapt the model for peptide-specific prediction, we then fine-tuned BindEvaluator on over 12,000 structurally validated peptide-protein pairs,^14,31^ achieving high precision in peptide-protein binding site prediction with an AUC of 0.97 (**Figure 1B, Supplementary Figures 1C**).

### moPPIt integrates BindEvaluator into MOG-DFM to design motif-specific peptide binders

Having trained BindEvaluator to accurately predict peptide-protein binding sites, we developed moPPIt to generate peptide binders that simultaneously optimize motif specificity and binding affinity from target protein sequences alone. moPPIt extends our recent Multi-Objective-Guided Discrete Flow Matching (MOG-DFM) framework,^20^ which steers a pretrained discrete flow matching generator toward Pareto-efficient trade-offs by modulating transition probabilities via rank-directional scoring and adaptive hypercone filtering. When previously applied to an unconditional peptide generator, PepDFM, MOG-DFM guidance simultaneously improved multiple competing therapeutically relevant properties.^20^ Here, we apply MOG-DFM to guide PepDFM using two complementary models: BindEvaluator to enforce user-defined motif engagement and a pre-trained, published affinity predictor^20^ to ensure sufficiently strong binding (**Figure 1C**).

To confirm that moPPIt’s underlying MOG-DFM framework can balance both motif specificity and binding affinity, we performed binder generation experiments targeting three different proteins: PDB entries 5AZ8, 7JVS, and MYC (**Table 1**). In an initial ablation study, we removed one or both objectives. The motif scores and affinity scores of the designed binders prove that omitting any single guidance causes a collapse in that property, while the other metric may modestly improve. In contrast, enabling both guidance signals yields the most balanced profiles across both objectives. Notably, this balanced performance holds across both structured proteins with known binders (5AZ8, 7JVS) and a disordered protein with more minimal experimental data (MYC), thus demonstrating moPPIt’s unique versatility.

**Table 1.**
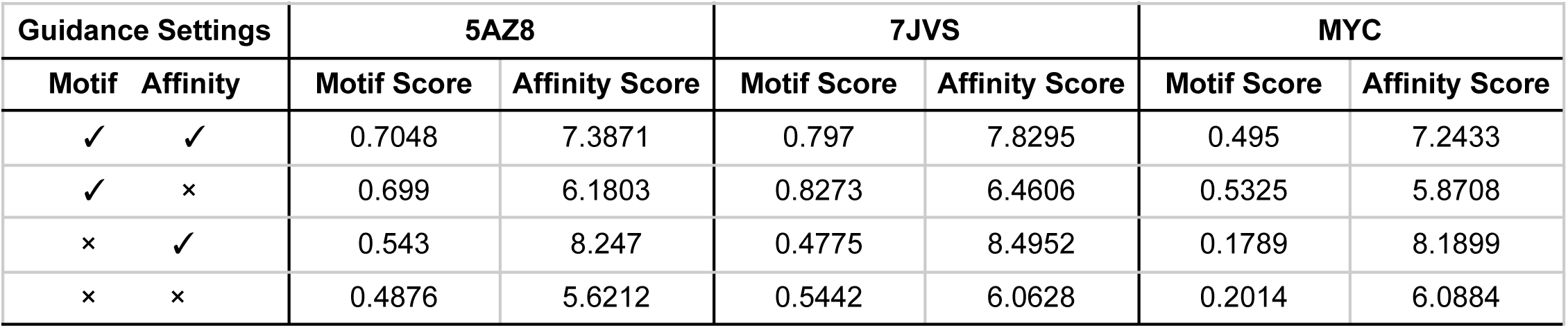
Ablation results for binder design targeting 5AZ8 (PDB ID), 7JVS (PDB ID), and MYC with different guidance settings. For each setting, 100 binders were designed with lengths of 11, 11, and 8, respectively. The average scores are displayed.

| Guidance Settings |  | 5AZ8 |  | 7JVS |  | MYC |  |
| --- | --- | --- | --- | --- | --- | --- | --- |
| Motif | Affinity | Motif Score | Affinity Score | Motif Score | Affinity Score | Motif Score | Affinity Score |
| ✓ | ✓ | 0.7048 | 7.3871 | 0.797 | 7.8295 | 0.495 | 7.2433 |
| ✓ | × | 0.699 | 6.1803 | 0.8273 | 6.4606 | 0.5325 | 5.8708 |
| × | ✓ | 0.543 | 8.247 | 0.4775 | 8.4952 | 0.1789 | 8.1899 |
| × | × | 0.4876 | 5.6212 | 0.5442 | 6.0628 | 0.2014 | 6.0884 |

### moPPIt generates epitope-specific binders to target proteins *in silico*

With moPPIt’s dual-objective optimization framework established, we tested its capabilities across three increasingly difficult tasks: targeting proteins with previously known binders, generating binders against novel epitopes within structured proteins, and designing binders against intrinsically disordered regions. We first sought to compare moPPIt-generated peptides with experimentally validated peptides not included within its training data. For 15 structured proteins within the PDB, we evaluated the quality of AlphaFold3 (AF3)-predicted peptide-protein complexes using ipTM scores (a measure of interface confidence), AutoDock VINA docking scores, and PeptiDerive residue-level binding free energy predictions.^32–34^. Overall, we observe that moPPIt-designed binders form peptide-protein complexes of comparable or superior quality to pre-existing binders across ipTM and VINA docking score (**Table 2**). Notably, all 15 designed peptides met stringent ipTM thresholds (within 0.05 of reference), while 12 of 15 exceeded the VINA score thresholds (set at 1.0 lower than that of the reference complex) (**Figure 1D, 1E**). Moreover, PeptiDerive analysis confirmed that high-energy residues localized to the target motifs in designed complexes, validating motif-specific engagement (**Supplementary Figures 2, 3**). As a whole, these *in silico* metrics indicate that moPPIt is capable of designing high quality, motif-specific peptides on structured targets without structural supervision.

**Table 2.**
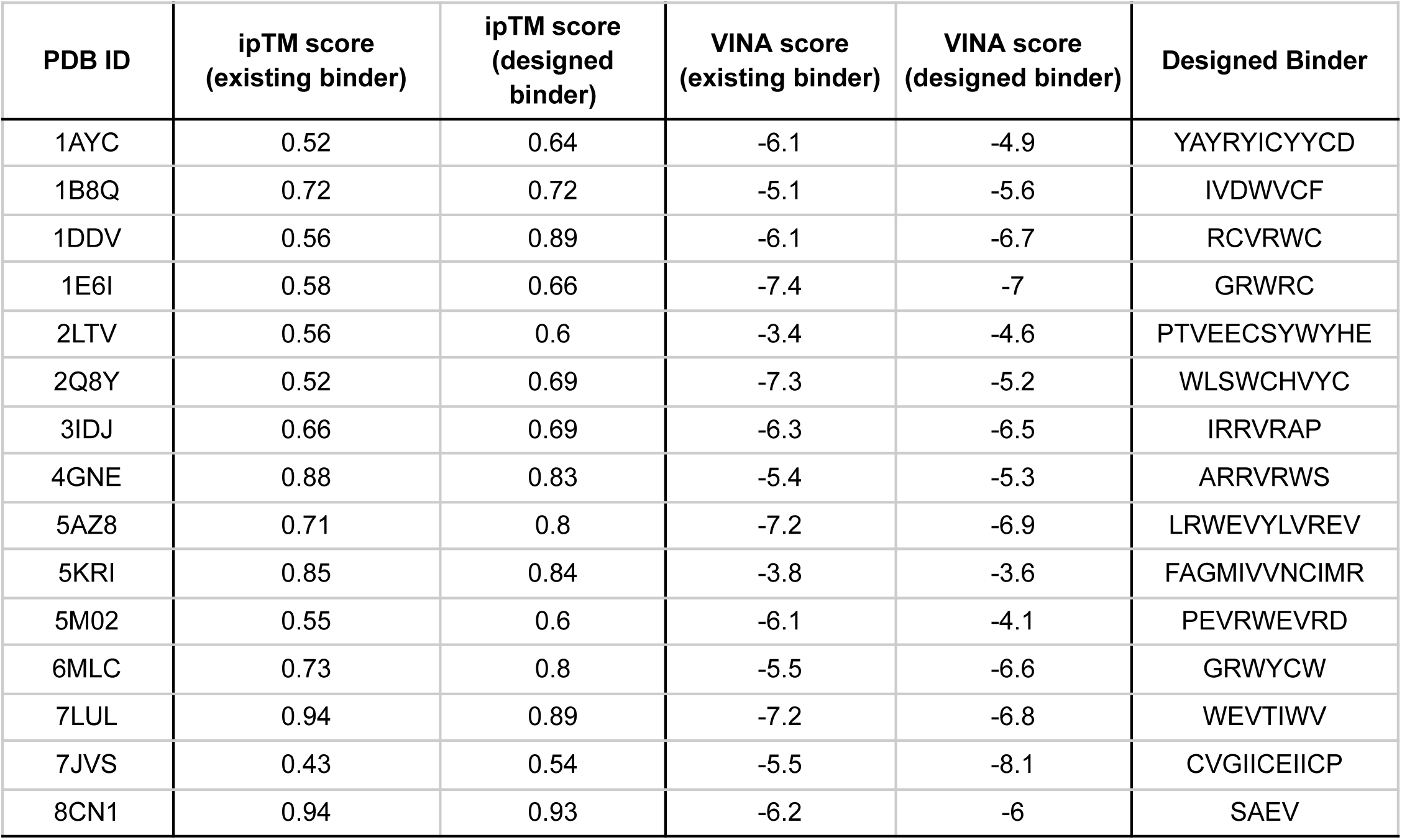
Comparison of ipTM for existing and designed peptide-protein complexes. The ipTM scores are calculated by AlphaFold3 for peptide-protein complexes using both existing peptides and peptides designed by the moPPIt algorithm. The designed binders for each protein are presented.

Next, we tasked moPPIt with designing binders for structured proteins lacking known peptide partners from diverse structural classes including kinases, phosphatases, deubiquitinases, and GPCRs. Targeting candidate binding sites identified using APBS electrostatic analysis, moPPIt successfully designed epitope-specific binders with high predicted affinities (mean VINA score of −8.2 kcal/mol) and motif-localized binding energies (**Table 3**). The latter is evidenced by PeptiDerive RIS analysis of predicted peptide-protein complexes, indicating localized amino acid interactions within or proximal to the targeted motif sequence (**Supplementary Figures 4, 5, 6, Supplementary Table 1**).^32^ Combined with 3D visualization of predicted structures, we demonstrate that even in cases where a peptide binder is unknown, moPPIt still yields peptides with high affinity for their target motifs.

**Table 3.**
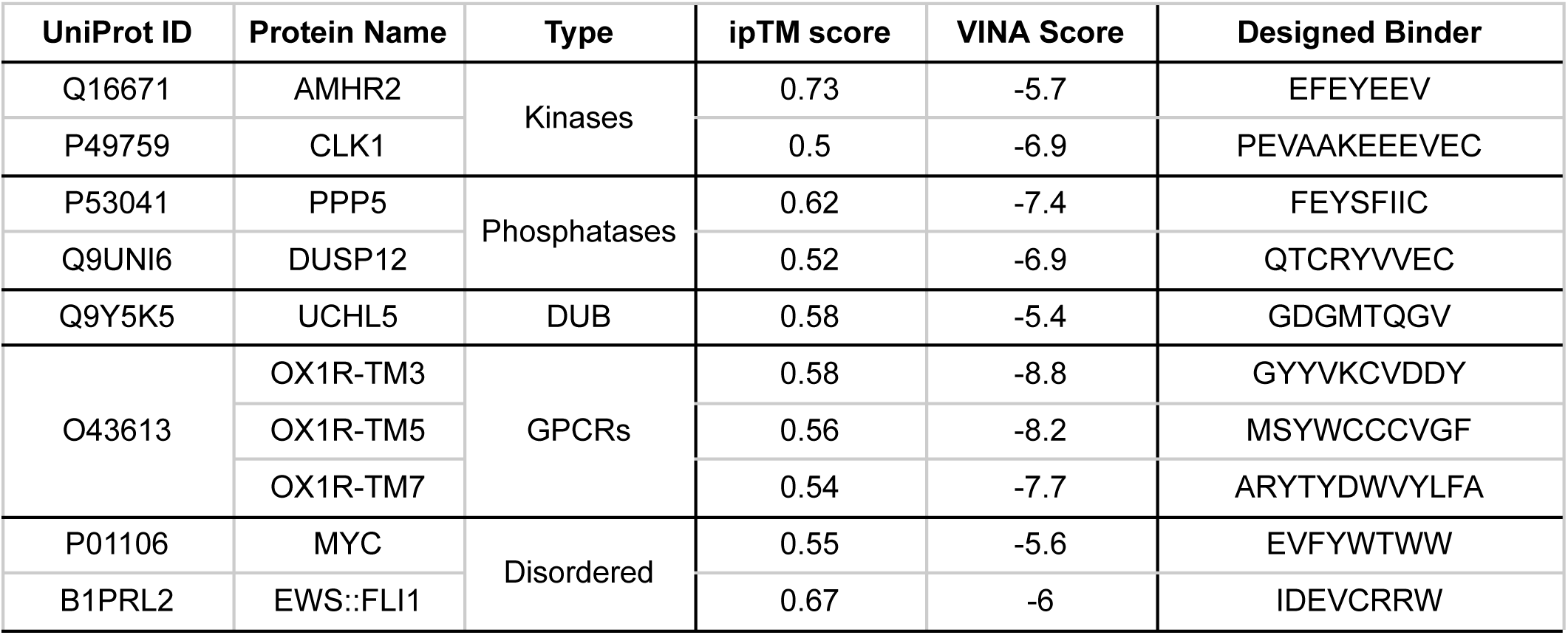
pTM and ipTM scores and VINA docking scores for designed binders targeting proteins without known binders. This table lists the pTM and ipTM scores for the complex structures of proteins with designed binders targeting proteins without known binders. The proteins are categorized by type, including kinases, phosphatases, and deubiquitinating enzymes (DUBs), GPCRs, and intrinsically disordered proteins. The designed binders and AutoDock VINA docking scores are provided alongside each protein.

Finally, we turned towards highly-challenging, “undruggable” intrinsically disordered proteins, selecting MYC and EWS::FLI1 to test moPPIt’s capabilities. Since these proteins lack stable 3D structures and sample transient conformational ensembles, they inherently limit the reliability of traditional structure-based design.^10,11^ Despite this ambiguity, AlphaFold3-predicted complexes of moPPIt-designed peptides suggest specific interaction with the targeted motifs, a finding further corroborated by favorable VINA docking scores and high PeptiDerive interface scores localized to the binding site (**Table 3, Supplementary Figure 7**). However, precisely because intrinsic disorder confounds static structural prediction, we posit that BindEvaluator’s motif score serves as the most fundamental validation metric in this regime. Unlike structural metrics that rely on potentially uncertain folded states, BindEvaluator confirms specificity purely from sequence data, demonstrating that moPPIt can effectively target the disordered proteome even when structural ground truth is elusive.

### moPPIt-generated binders show motif and domain-specificity *in vitro*

We next turned to validating moPPIt experimentally via design of peptides to numerous, clinically-relevant targets (**Supplementary Table 2**). First, in order to ensure moPPIt’s capacity for domain specific binding, we designed peptides against two distinct domains of neural cell adhesion molecule 1 (NCAM1) (**Figure 2A**), a key marker of acute myeloid leukemia.^35^ NCAM1 is a seven-domain glycoprotein composed of five immunoglobulin (IgG) domains followed by two FN3 domains, each with distinct structures and cell-signaling function (**Supplementary Table 3**).^36^ Using moPPIt, we generated 100 10-mer peptides specifically targeting the second FN3 domain (residues 611-706) of NCAM1. Of these sequences, the top five (based on motif and affinity score) were selected for *in vitro* validation and expressed in *E. coli* as C-terminal fusions to a SUMO-tag protein (**Supplementary Table 2**). Proper engagement of the target residues within the FN3 region was also confirmed using AlphaFold3 structure prediction (**Figure 2B**).

**Figure 2:**
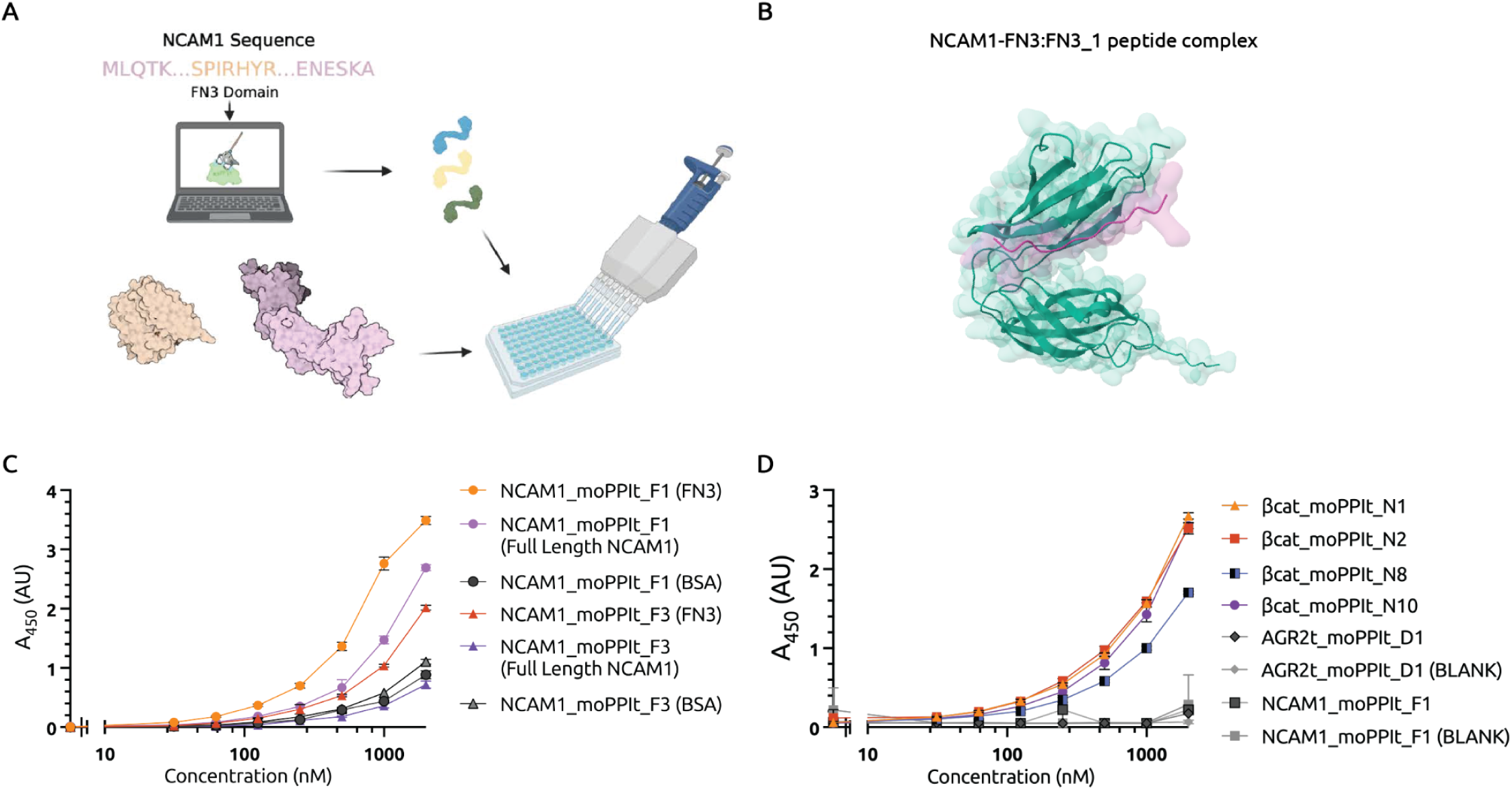
*in vitro* validation of NCAM1-FN3 and β-catenin intrinsically disordered region (IDR) peptide binders. **(A) Schematic of moPPIt workflow for *in vitro* validation.** Target protein sequence and binding motif (residue numbers) are input to generate optimized peptide sequences, which are then expressed in E. coli as C-terminal fusions to a SUMO-tag, then subsequently validated via ELISA. **(B)** AlphaFold3-predicted structure of NCAM1_moPPIt_F1 peptide (purple) bound to the FN3 domain of NCAM1 (green), demonstrating predicted engagement with the target epitope (residues 611-706). **(C) Validation of domain-specific binding of NCAM1-targeting peptides.** Five moPPIt-generated peptides targeting the second FN3 domain of NCAM1 were expressed as SUMO-tagged fusions and tested by ELISA against isolated NCAM1-FN3 (domains 1-2; residues 509-706), full-length NCAM1, and BSA controls. Two peptides (NCAM1_moPPIt_F1, NCAM1_moPPIt_F3) showed low nanomolar binding to isolated NCAM1-FN3 with minimal BSA background. NCAM1_moPPIt_F1 retained strong binding to full-length NCAM1, confirming specific engagement with the FN3 domain within the intact protein. **(D) Validation of binding to β-catenin IDR-targeting peptides.** Four moPPIt-designed peptides targeting the N-terminal IDR of β-catenin (residues 1-150) and two off-target control proteins (AGR2t and NCAM1-FN3) were tested against full-length β-catenin by ELISA. All four candidates showed robust binding to β-catenin compared to peptides designed to target AGR2t and NCAM1-FN3 (AGR2t_moPPIt_D1, NCAM1_moPPIt_F1), demonstrating that moPPIt can generate favorable binders to conformationally flexible regions inaccessible to structure-based design methods.

Two of the five peptides (NCAM1_moPPIt_F1, NCAM1_moPPIt_F3) demonstrated domain-specific interaction, both of which bound to isolated NCAM1-FN3 with observable EC50 values in the low nanomolar range. For both peptides, off-target interactions with BSA controls were minimal (**Figure 2C**). Critically, NCAM1_moPPIt_F1 retained strong binding to full-length NCAM1, confirming engagement with the FN3 domain rather than potential artifacts from the isolated construct. This validates moPPIt’s capability to target specific regions within multi-domain proteins.

We next challenged moPPIt with targeting IDRs within a pathogenically relevant target. The N-terminal IDR of β-catenin (residues 1-150) harbors potential disease-driving mutations and contains the β-TrCP1 E3-ligase recognition sequence required for proteasomal degradation (**Supplementary Table 3**).^37^ Binding either to subsequences containing these potential mutants or the β-TrCP1 degron itself could prove useful in modulating β-catenin function or degradation kinetics at the cellular level in diseases such as hepatocellular carcinoma.^38^ From 100 moPPIt-generated sequences targeting this IDR, ten were expressed and preliminarily screened by ELISA (**Supplementary Table 2**). The top four candidate peptides, selected by signal intensity and signal-to-noise ratio, all showed robust binding to full-length β-catenin compared to other moPPIt-derived peptides targeting unrelated protein controls (AGR2t, NCAM1-FN3) (**Figure 2D**). These results demonstrate that moPPIt can generate specific binders to conformationally flexible regions inaccessible to structure-based design methods.

### moPPIt peptides bind GM-CSFRα and block macrophage activation

To move from biophysical binding to functional disease modulation, we sought to design inhibitors against the granulocyte-macrophage colony-stimulating factor (GM-CSF) receptor. GM-CSF is a cytokine classically implicated in monocyte-to-macrophage differentiation and activation. Emerging evidence indicates that GM-CSF signaling promotes the generation of profibrotic macrophage populations with extracellular matrix (ECM) remodeling capacity in diverse tissues.^26,39–41^ In light of these findings, we used moPPIt to design peptide inhibitors to disrupt GM-CSF-driven profibrotic macrophage polarization by targeting the receptor alpha subunit (CSF2Rɑ), which mediates ligand binding (**Supplementary Tables 2-3**). Three peptide classes were generated: (i) peptides engaging the CSF2Rɑ ligand-binding pocket, (ii) peptides directed against the conserved WSxWS motif, and (iii) motif-independent binders. In addition to affinity- and motif-score-based guidance, these peptides were generated via multi-objective guidance on high solubility and low hemolysis scores from our previously-published predictors^20^ to ensure successful synthesis and low peptide-level toxicity in culture.

Freshly isolated human peripheral blood monocytes from buffy coats were pre-incubated with individual peptides (5 µM) for 2 hours and subsequently cultured in the continuous presence of GM-CSF (50 ng/ml) for 72 hours, with daily medium replacement. Monocyte-to-macrophage differentiation was quantified by cell size measurements from brightfield microscopy images as a surrogate for macrophage maturation (see *Methods*). Across the panel, almost all moPPIt-designed peptides reduced GM-CSF-induced differentiation, with P3, P5, and MF3 demonstrating the greatest inhibitory effect, showing macrophage area z-scores comparable to untreated negative controls (**Figure 3A**). Representative brightfield microscopy images further confirm this functional blockade; stimulated cells treated with moPPIt inhibitors failed to undergo the characteristic transition from small, rounded monocytes to the large, spread morphology seen in the GM-CSF-only positive control (**Figure 3B**). Notably, this potent inhibition across varied receptor motifs demonstrates that moPPIt can successfully disrupt complex signaling assemblies by precisely targeting distinct functional epitopes.

**Figure 3:**
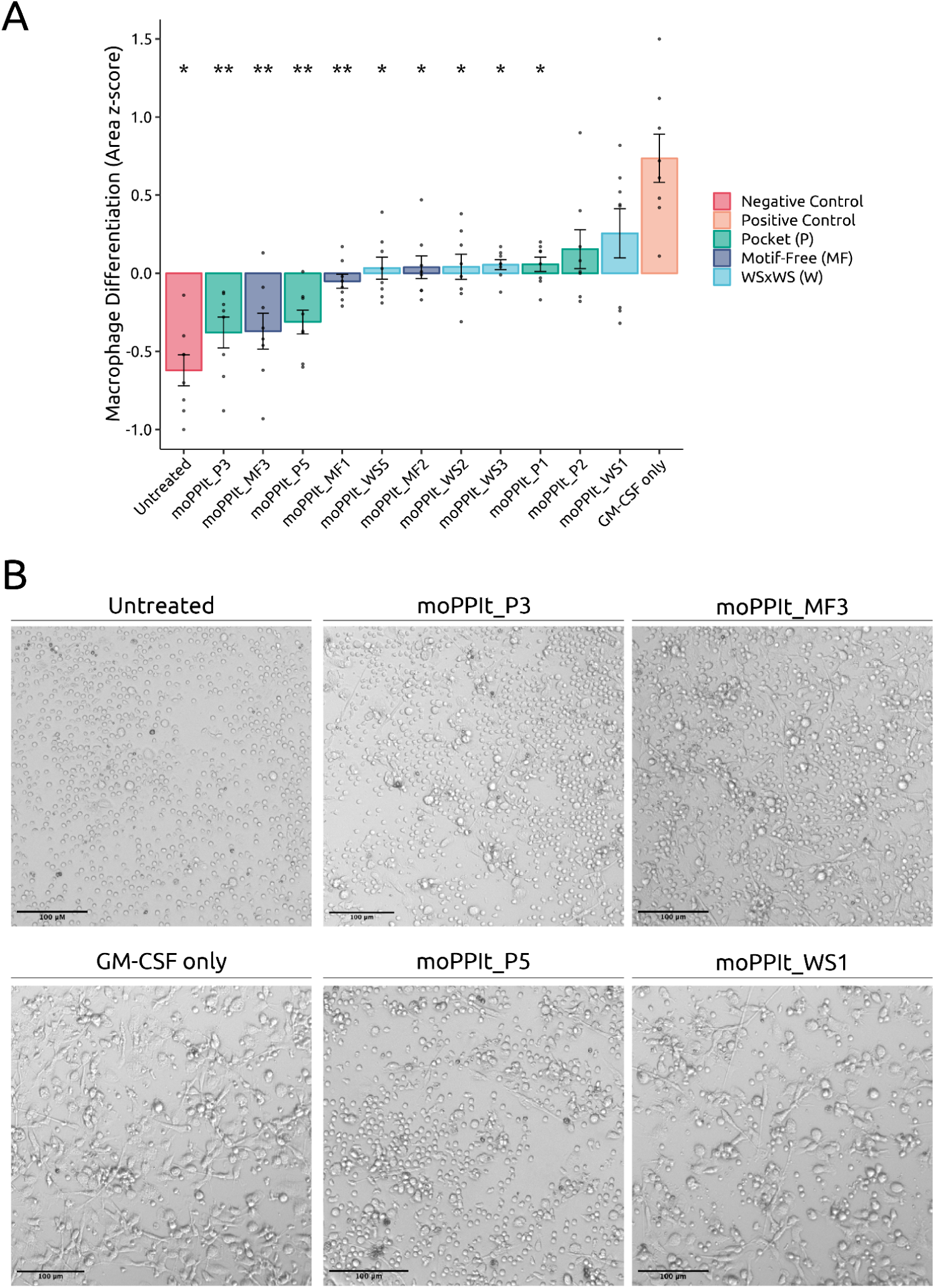
Inhibition of macrophage differentiation using moPPIt-generated peptides targeting GM-CSF receptor α subunit. **(A) Bar plot showing the mean z-scored cell areas across conditions.** Monocyte-to-macrophage differentiation was quantified by cell area (z-scored by donor) following 72-hour treatment with GM-CSF (50 ng/ml) in the presence or absence of moPPIt peptides (5 µM). Freshly isolated primary human monocytes were pre-incubated with peptides for 2 hours prior to GM-CSF stimulation, with media refreshed daily. Larger cell areas indicate enhanced macrophage differentiation. Three classes of peptides were designed: Pocket (P), targeting the GM-CSFα ligand-binding pocket; WSxWS (WS), targeting the conserved WSxWS activation motif; and Motif-Free (MF), generated without structural constraints. Compared to GM-CSF alone (positive control), all moPPIt-designed peptides strongly reduced macrophage differentiation, with P3, P5, and MF3 exhibiting near-complete inhibition. Statistical comparisons performed using pairwise Wilcoxon rank-sum tests with Bonferroni correction. *p < 0.05, **p < 0.01, ns = not significant (n = 8 donors). **(B) Representative brightfield microscopy images (100 µm scale bars) showing cell morphology across treatment conditions.** Note the transition from small, rounded monocytes (Untreated) to large, spread macrophages (GM-CSF only), which is attenuated by peptide treatment.

### moPPIt-generated binder-based chimeric antigen receptor (CAR) activates human regulatory T cells with suppressive function

Regenerative medicine holds promise to provide treatments for many diseases by replacing damaged cells, tissues, and organs with new ones generated from human induced pluripotent stem cells (hiPSCs). A major hurdle to fulfill this vision is immunological rejection of genetically mismatched tissues by recipients.^42^ Regulatory T cells (Tregs) are a subset of immune cells endowed with immunosuppressive properties.^43^ Utilizing them to prevent the rejection of transplanted tissues requires employing Tregs that specifically recognize those tissues. Chimeric antigen receptors (CARs) are synthetic immune receptors containing an antigen binding domain and an intracellular T cell signaling domain that can be used to define Treg specificity.^43,44^ We have recently shown that CAR Tregs can traffic to and protect transplanted hiPSC-derived beta cells expressing a cell surface ligand recognized by the CAR from immune rejection.^44^ To generalize this approach, we first created a synthetic unique cell surface ligand that is biologically and immunologically inert by deleting the endoplasmic retention motif (KTEL) and mutating the CXXS thioredoxin binding motif (C81S) of anterior gradient protein 2 (AGR2)^45^ and fusing it to the transmembrane domain of platelet derived growth factor receptor (PDGFR) (**Supplementary Table 3**). K562 cells, a human cell line that lacks human leukocyte antigen (HLA) surface expression and hence cannot be recognized by T cells, transduced with this construct, which we called AGR2t, stably expressed the protein on the surface (**Supplementary Figure 8A**). The antigen binding domain of a CAR is most commonly a ~250 amino acid antibody-derived single chain variable fragment (scFv). This design represents a major bottleneck in developing new CARs, as it necessitates a costly, laborious, and long process to generate antibodies, format them into scFvs, and test if CARs incorporating these scFvs are expressed on the surface of T cells and lead to T cell activation upon binding to the desired target.^46,47^ To address these limitations and generate a synthetic CAR-ligand pair, we employed moPPIt to generate 12 amino acid-long peptides binding specifically to the extracellular portion of AGR2t. Peptide sequences were initially evaluated based on motif score, affinity score, and VINA docking scores (**Figure 4A**). Top scorers in each category were tested for binding to a subfragment (amino acids 21-144) of AGR2 protein using enzyme linked immunosorbent assay (ELISA), with bovine serum albumin (BSA) as a negative control (**Figure 4B**). Most peptides tested specifically bound to AGR2 protein (**Figure 4B**).

**Figure 4.**
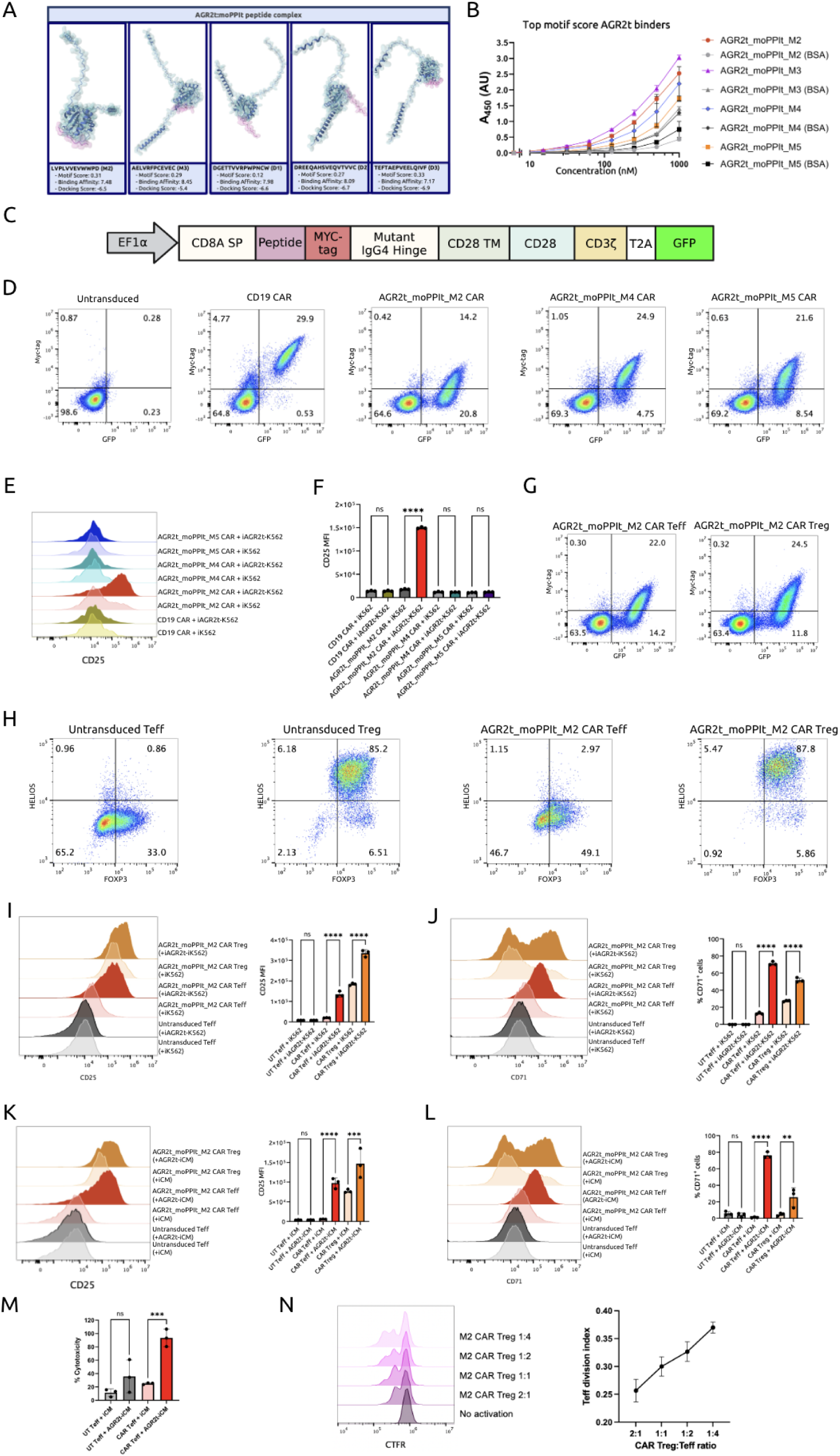
*De novo* generation of synthetic ligand-specific chimeric antigen receptor effector and regulatory T cells using moPPIt. **(A)** moPPIt-designed peptides (pink) docked to AGR2t (green) via AlphaFold3 and scored via VINA docking. Docking scores are provided in kcal/mol. **(B)** Sandwich ELISA with immobilized AGR2t treated with increasing concentrations of specified peptides. **(C)** Chimeric antigen receptor (CAR) schematic featuring moPPIt generated AGR2t-specific peptide binder sequences linked to a Myc-tag and a mutant IgG4 hinge sequence, a CD28-CD3ζ signaling domain, and a 2A peptide-linked GFP reporter gene. **(D)** CAR surface expression (Myc-tag) and GFP reporter expression in lentivirus transduced human CD4+ T cells. UT, untransduced. **(E-F)** Surface expression of T cell activation marker CD25 on CD4+ T cells expressing CARs with different moPPIt generated AGR2t-specific binder sequences upon 48h co-incubation with irradiated (i) AGR2t-expressing K562 cells. Representative histogram on the left and summary data on the right. MFI, median fluorescence intensity. **(G)** AGR2t_M2_CAR surface expression (Myc-tag) and GFP reporter expression in lentivirus transduced human Teff cells and Tregs. **(H)** Expression of Treg lineage transcription factors FOXP3 and HELIOS in UT and AGR2t_M2_CAR Teff cells and Tregs. i. Surface expression of CD25 on AGR2t_M2_CAR Teff cell and Tregs upon 48h co-incubation with irradiated AGR2t-expressing K562 cells. Representative histogram on the left and summary data on the right. **(I)** Surface expression of T cell activation marker CD71 on AGR2t M2 CAR Teff cell and Tregs upon 48 h co-incubation with irradiated AGR2t-expressing K562 cells. Representative histogram on the left and summary data on the right. **(J)** Surface expression of AGR2t in human induced pluripotent stem cell-derived cardiomyocytes (iCM). Shaded histogram represents untransduced iCMs and line histogram represents AGR2t lentivirus transduced iCMs. **(K)** Surface expression of CD25 on AGR2t_M2_CAR Teff cell and Tregs upon 48 h co-incubation with irradiated AGR2t-expressing iCMs. Representative histogram on the left and summary data on the right. **(L)** Surface expression of CD71 on AGR2t M2 CAR Teff cell and Tregs upon 48 h co-incubation with irradiated AGR2t-expressing iCMs. Representative histogram on the left and summary data on the right. **(M)** Cytotoxic activity of AGR2t M2 CAR Teff cells towards AGR2t-expressing iCMs. **(N)** Suppressive activity of AGR2t_M2 _CAR Tregs activated by irradiated AGR2t-K562 cells towards polyclonally activated Teff cells, as measured by inhibition of Teff cell proliferation. Representative histograms on the left and summary data for Teff cell division index on the right. Points represent mean and standard deviation. Bars in e, i, j, l, m, and n represent mean and standard deviation and statistical significance was performed using one-way ANOVA with pairwise comparisons, n = 3 replicates. **, p < 0.01; ***, p < 0.001; ****, p < 0.0001; ns, not significant.

Based on moPPIt scores and ELISA biochemical binding results, we prioritized peptides AGR2t_moPPIt_M2, AGR2t_moPPIt_M4, and AGR2t_moPPIt_M5 for functional testing using the CAR assay. Because moPPIt peptides are 20 times shorter than a traditional scFv, we reasoned that they require a longer hinge domain to function in the context of a CAR. We thus generated CAR constructs where the AGR2t_moPPIt_M2, AGR2t_moPPIt_M4, or AGR2t_moPPIt_M5 peptide sequence was followed by a Myc-tag to detect surface expression by flow cytometry and a long mutant IgG4 hinge, followed by the commonly employed CD28 transmembrane domain and CD28-CD3ζ signaling domain^44,48^ (**Figure 4C**). A GFP reporter gene linked to the CAR gene by a 2A peptide was included to allow assessment of T cell transduction independently of CAR surface expression (**Figure 4C**). We purified CD4^+^ T cells from human peripheral blood with lentivirus encoding for each of these new CARs, as well as with a published CD19CAR-2A-GFP construct^49^ as a negative control.

All CARs were expressed on the surface of human CD4^+^ T cells, as assessed by the presence of Myc-tag^+^GFP^+^ cells by flow cytometry (**Figure 4D**). To evaluate CAR function, we co-incubated CAR CD4^+^ T cells with irradiated AGR2t-expressing K562 cells. AGR2t_moPPIt_M2 CAR CD4^+^ T cells, but not AGR2t_moPPIt_M4 CAR, AGR2t_moPPIt_M5 CAR, or control CD19 CAR CD4^+^ T cells, were activated by AGR2t K562 cells, as assessed by upregulation of the T cell activation marker CD25 (**Figure 4E**). Next, we sought to determine whether the AGR2t_moPPIt_M2 CAR was also functional in human Tregs using our established protocols.^50^ We sorted CD4^+^CD25^high^CD127^low^ Tregs and CD4^+^CD25^low^CD127^high^ effector T cells (Teff) by fluorescence-activated cell sorting (FACS) (**Figure 4F**) and transduced each subset with AGR2t_moPPIt_M2 CAR. In line with our findings with bulk CD4+ T cells, both Tregs and Teff cells expressed AGR2t_moPPIt_M2 CAR on the surface. Importantly, AGR2t_moPPIt_M2 CAR Tregs maintained expression of the Treg lineage transcription factors FOXP3 and HELIOS (**Figure 4H**), indicating a stable suppressive Treg phenotype.^43^ Moreover, AGR2t_moPPIt_M2 CAR Tregs and AGR2t_moPPIt_M2 CAR Teff cells were activated by AGR2t-K562 cells, upregulating surface expression of CD25 (**Figure 4I**) and CD71 (**Figure 4J**), an additional T cell activation marker.^49^ Of note, AGR2t_moPPIt_M2 CAR Treg basal levels of CD25 were almost as high as those of activated AGR2t_moPPIt_M2 CAR Teff cells, further indicative of a suppressive Treg phenotype.^43^

Having established that AGR2t_moPPIt_M2 CAR can effectively elicit activation in Tregs and Teff cells without any deleterious effect upon recognition of AGR2t-K562 cells, we introduced AGR2t in a clinically relevant cell type: human induced pluripotent stem cell-derived cardiomyocytes (iCMs). Cardiovascular disease (CVD) is the number one cause of death in the world. Given the negligible endogenous regenerative potential of the heart, CVD stands to benefit from regenerative medicine approaches, such as iCM-based cell therapy. Protecting iCMs with CAR Tregs could allow the use of off-the-shelf iCMs for cardiac repair in any patient. Using our established protocols,^51,52^ we differentiated human induced pluripotent stem cells into iCMs and transduced them with AGR2t. After confirming AGR2t surface expression in transduced iCMs, we co-incubated either unmodified or AGR2t-iCMs with AGR2t_moPPIt_M2 CAR Tregs or AGR2t_moPPIt_M2 CAR Teff cells. In line with our findings with AGR2t-K562 target cells, AGR2t_moPPIt_M2 CAR Tregs and AGR2t_moPPIt_M2 CAR Teff cells upregulated CD25 (**Figure 4K**) and CD71 (**Figure 4L**) specifically in the presence of AGR2t-iCMs, demonstrating M2 CAR activation by a clinically relevant cell type. Moreover, AGR2t_moPPIt_M2 CAR Teff cells were cytotoxic specifically towards AGR2t-iCMs (**Figure 4M**) and AGR2t_moPPIt_M2 CAR Tregs suppressed Teff proliferation (**Figure 4N**), establishing that the AGR2t_moPPIt_M2 CAR is functional in human Teff cells and in Tregs.

## Discussion

The design of highly specific and affine peptide binders, particularly for targets lacking well-defined structural pockets or those possessing IDRs, has long represented a bottleneck in therapeutic development.^1^ In this work, we presented **moPPIt**, a purely sequence-based approach that addresses this challenge by enabling the *de novo* design of motif-specific peptide binders, independent of structural representations. By leveraging a discrete flow matching-based multi-objective optimization framework,^20^ moPPIt generates peptides that engage user-defined epitopes across a broad spectrum of targets, including those with structured and conformationally flexible motifs, and further exhibit clinically-relevant effects for human macrophage polarization disruption and CAR Treg cell activation, enabling therapeutically-ready applications.

Given these comprehensive results, we anticipate that moPPIt will be effective across diverse protein classes. While out of scope here, in a future study, we will experimentally evaluate moPPIt alongside structure-based methods such as RFDiffusion and BindCraft,^7,8^ evaluating performance on both ordered and disordered regions. A critical component of this validation will be the conversion of moPPIt-designed peptides into functional effectors. We plan to deploy these binders within chimeric proteome editing architectures, specifically ubiquibodies (uAbs) and deubiquibodies (duAbs) to mediate targeted protein degradation and stabilization studies.^13,53–55^ Crucially, to enable the deployment of standalone peptide therapeutics, we will expand the moPPIt framework to explicitly optimize additional developability metrics, such as protease resistance, immunogenicity, and half-life, directly during the design phase. We further plan to extend these capabilities to non-canonical and cyclic peptides, building on our work with PepTune, TR2-D2, and our new rectified discrete flow matching framework, AReUReDi.^17,18,22^ Such multi-objective guidance ensures that generated candidates possess not only high affinity and specificity but also the pharmacokinetic properties required for clinical translation.

Furthermore, the motif-specific nature of our approach suggests promising applications in developing binders with exquisite selectivity, including the ability to discriminate between pathogenic mutants and wild-type variants,^56^ or to target specific post-translational modification sites.^57^ Importantly, the capacity to target specific epitopes is particularly valuable for interrogating viral proteins, such as those of SARS-CoV-2 and future pandemic threats,^14,58^ as well as to induce immune tolerance to specific disease-causing antigens in autoimmune disease and organ transplant rejection^14,5843^ moPPIt enables the design of binders against highly conserved regions that are less prone to escape mutations. Collectively, these capabilities highlight moPPIt as a powerful tool for both detection and therapeutic applications, enabling the precise modulation of protein function in disease contexts.

## Methods

### Discrete Flow Matching

In the discrete setting, we consider data *x* = (*x*_1_,…,*x_d_*) taking values in a finite state space *S* = *𝒯^d^*, where *𝒯* = [*K*] = {1,2,…,*K*} is called the vocabulary. We model a continuous-time Markov chain (CTMC) {*x_t_*}*_t_*_∈[0,1]_ whose time-dependent transition rates *u_t_*(*y,x*) transport the probability mass from an initial distribution *p*_0_ to a target distribution *p*_1_.^59^ The marginal probability at time *t* is denoted *p_t_*(*x*), and its evolution is governed by the Kolmogorov forward equation:

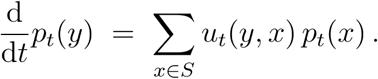

The learnable velocity field *u_t_*(*y,x*) is defined as the sum of factorized velocities:

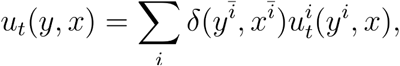

where *ī* = (1,…,*i*−1,*i*+1,…,*d*) denotes all indices excluding *i*. The rate conditions for factorized velocities 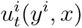 are required per dimension *i* ∈ [*d*]:

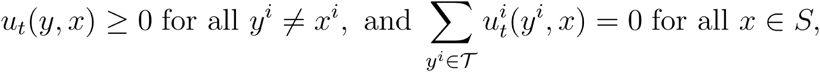

so that for small *h* > 0, the one-step kernel

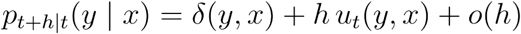

remains a proper probability mass function.

In practice, we can further parameterize the velocity field using a mixture path. Specifically, a mixture path is defined with scheduler *k_t_* ∈ [0,1] so that each coordinate 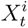 equals 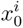 or 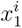 with probabilities 1 − *k_t_* and *k_t_*, respectively. The mixture marginal velocity is then obtained by averaging the conditional rates over the posterior of (*x*_0_. *x*_1_) given *X_t_* = *x*, yielding

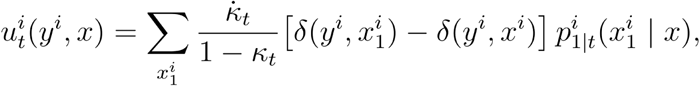

where *k̇_t_* denotes the time derivative of *k_t_*.

### moPPIt Formulation

moPPIt operates under the same setting as discrete flow matching described in the previous section. At its core, it leverages a pretrained discrete flow matching model, PepDFM, that defines a CTMC with a factorized velocity field 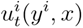, which transports probability mass from an initial distribution *p*_0_ to the target distribution over plausible peptide sequences via mixture path parametrization. In addition, moPPIt uses two pretrained score models, the affinity predictor *s*_1_ and BindEvaluator *s*_2_, that assigns objective scores to any peptide sequence. The affinity predictor *s*_1_ calculates the affinity score based on the peptide-protein pair, while *s*_2_ predicts the motif score given a target protein sequence, a peptide binder sequence, and target motifs. Specifically, motif score is calculated as the average probability of each target motif residue participating in binding, as predicted by BindEvaluator:

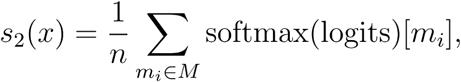

where *M* represents the target motifs.

We aim to generate novel sequences *x*_1_ ∈ *𝒮* whose objective vectors (*s*_1_(*x*_1_), *s*_2_(*x*_1_)) lie near the Pareto front (not guaranteed to be Pareto optimal)

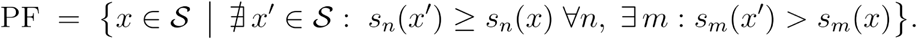

To achieve this, moPPIt applies the MOG-DFM framework that guides the CTMC sampling dynamics of PepDFM using multi-objective transition scores, steering the generative process toward Pareto-efficient regions of the state space.

moPPIt begins by initializing the generative process at time *t* = 0 by sampling an initial sequence *x*_0_ uniformly from the discrete state space *𝒮* = [*K*]*^d^*. To steer the generation towards diverse Pareto-efficient solutions, we introduce a set of *M* weight vectors 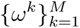, where w ∈ ℝ^2^, that uniformly cover the 2-dimensional Pareto front. Intuitively, each *ω* encodes a particular trade-off among both objectives, so sampling different *ω* promotes exploration of distinct regions of the Pareto front. We construct these vectors via the Das-Dennis simplex lattice with *H* subdivisions, yielding components

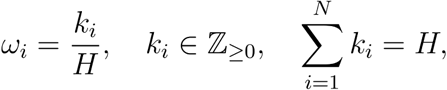

A single *ω* is then sampled randomly to define the optimization direction toward the Pareto front for the current run. Once initialized, moPPIt performs Step 1 (Guided Transition Scoring), Step 2 (Adaptive Hypercone Filtering), and Step 3 (Euler Sampling) over *T* iterations to generate a final sequence *x*_1_ whose score vectors have been steered close to the Pareto front, with both objectives optimized. For detailed formulations of these steps, please refer to Chen, et al.^20^

### Dataset Preparation

The training data for BindEvaluator was curated from the PPIRef dataset, a large and non-redundant database of PPIs with annotated binding sites.^30^ To augment the dataset, additional entries were generated by reversing the roles of the target and binder sequences for each original entry. Proteins exceeding 500 amino acids were removed due to GPU constraints. After removing all duplicates, the final dataset comprised 510,804 triplets, each containing a target sequence, a binder sequence, and binding motifs. This dataset was split at a 60/20/20 ratio into a training set, validation set, and test set.

The peptide-protein interaction data for fine-tuning BindEvaluator was curated from the PepNN and BioLip2 databases.^31,60^ Specifically, 3022 PepNN and 9251 BioLip2 non-redundant triplets for peptide-protein binding were collected. Proteins longer than 500 amino acids and peptides longer than 25 amino acids were removed. The dataset was split at an 80/10/10 ratio into a training set, validation set, and test set.

We collected 1,781 binding affinity data for classifier training from the PepLand and PeptideBERT datasets.^61,62^ All sequences taken are wild-type L-amino acids and are tokenized and represented by ESM-2 protein language model.^29^

### BindEvaluator Model Architecture

The generation algorithm is based on the BindEvaluator model. As shown in Figure 1B, BindEvaluator takes a binder sequence and a target sequence as inputs to predict the binding residues on the target protein. The design of this model draws inspiration from the architectures of PepNN and Pseq2Sites, which have demonstrated effectiveness in similar tasks.^29,31,63^

Both binder and target sequences are first passed into the pretrained ESM-2-650M model to obtain their embeddings.^29^ For the target sequence, a dilated CNN module captures the local features of adjacent residues. Specifically, the module is composed of three stacked CNN blocks with different dilation rates (1, 2, and 3) to extract hierarchical features. Each block consists of three convolutional layers with different kernel widths (3, 5, and 7) to cover different receptive field sizes, accommodating different binding site sizes. Padding is added to each convolutional layer to maintain consistent output and input sizes. Since the focus is to identify binding residues for the target protein, the dilated CNN module is applied only to the target sequence. Given that no binding motifs in the training set contain more than 23 continuous residues, the dilated CNN module is sufficient to capture the binding region features.

The processed embeddings are then passed through multi-head attention modules to capture global dependencies for each residue. In the reciprocal attention modules, the target and binder sequence representations are integrated to capture binder-target interaction information. Specifically, in these modules, the binder representations are projected into a key matrix *K* and a query matrix *Q*, while the target representations are projected into a value matrix *V*, and vice versa. The reciprocal attention is formulated as follows:

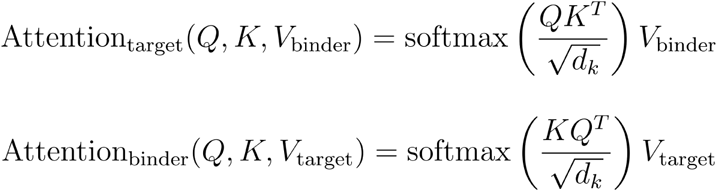

where *d_k_* is the model dimension.

Following several layers of dilated CNN and attention modules, the resulting target sequence representation encapsulates the binder-target binding information. Finally, this representation is processed by feed-forward layers and linear layers to predict the binding sites.

### BindEvaluator Training and Fine-Tuning

BindEvaluator is first trained on a protein–protein interaction (PPI) dataset and then fine-tuned using peptide-protein binding data. During training and fine-tuning, we use the same model architecture (Fig. 1A). The weights of ESM-2-650M are fixed throughout, and all other parameters remain trainable.

#### Loss Function

To accurately capture the intrinsic distribution of binding residues, the loss function *L* is designed to be the sum of the Binary Cross-Entropy (BCE) loss and the Kullback-Leibler (KL) divergence between the predicted and the true binding motifs. Specifically, letting *ŷ* be the predicted binding motifs and *y* be the true binding motifs, the loss function is defined as:

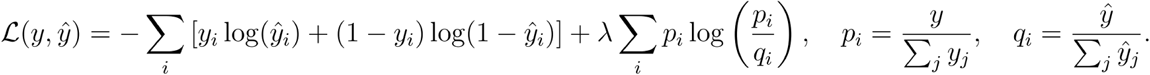

Here, *λ* is a hyperparameter that balances the contribution of the KL divergence to the total loss. During training, *λ* is set to 0.1, while during fine-tuning, *λ* is set to 1.

#### Comparison of BindEvaluator variants (with/without dilated CNN)

We first trained BindEvaluator without dilated CNN modules on a large protein-protein interaction dataset. During training, we observed a consistent decline in the validation loss, which indicates stable and effective learning (**Supplementary Figure 1A**). The steady decrease in binary cross entropy (BCE) loss and Kullback-Leibler (KL) divergence loss suggested that the model improves in distinguishing between binding and non-binding residues and in understanding the fundamental distribution of binding sites. We then trained BindEvaluator with dilated CNN modules on the same dataset. Both models, with and without dilated CNN modules demonstrated similar declining trends in their loss curves, indicating effective learning (**Supplementary Figure 1B**). Notably, the total loss continued to decrease even in the final training epochs, suggesting that the BindEvaluator with dilated CNN modules was more adept at learning subtle features, leading to better performance. During fine-tuning on the peptide-protein interaction dataset, we observed validation loss decreasing steadily, indicating a steady improvement in binding site prediction abilities (**Supplementary Figure 1C**).

#### Implementation

BindEvaluator was trained on a 6xA6000 NVIDIA RTX GPU system with 48 GB of VRAM each for 30 epochs. The batch size was set to 32, with a learning rate of 1e-3, a dropout rate of 0.3, and a gradient clipping value of 0.5. The AdamW optimizer was used with weight decay. Fine-tuning was performed on the same six GPUs for 30 epochs, with an increased dropout rate of 0.5. The batch size, learning rate, gradient clipping, and optimizer settings were identical to those used during training.

### Affinity Predictor

We previously developed an unpooled reciprocal attention transformer model to predict protein-peptide binding affinity,^20^ leveraging latent representations from the ESM-2-650M protein language model.^29^ Instead of relying on pooled representations, the model retains unpooled token-level embeddings from ESM-2-650M, which are passed through convolutional layers followed by cross-attention layers. The binding affinity data was split into a 0.8/0.2 ratio, maintaining similar affinity score distributions across splits. We used OPTUNA for hyperparameter optimization, tracing validation correlation scores.^64^ The final model was trained for 50 epochs with a learning rate of 3.84e-5, a dropout rate of 0.15, 3 initial CNN kernel layers (dimension 384), 4 cross-attention layers (dimension 2048), and a shared prediction head (dimension 1024) in the end. The classifier reached 0.64 Spearman’s correlation score on validation data.

### moPPIt Sampling

Hyperparameters related to MOG-DFM were set as follows: The number of divisions used in generating weight vectors, num_div, was set to 64, *λ* to 1.0, β to 1.0, *α_r_* to 0.5, *τ* to 0.3, *η* to 1.0, Φ*_init_* to 45°, Φ*_min_* to 15°, Φ*_max_* to 75°. The total sampling step *T* was 100. The importance vector was set to [20, 1], corresponding to motif score and affinity score, respectively.

### Expression and purification of SUMO-peptide constructs

Peptides of interest were cloned into a pET-24a+ (Novagen) expression vector containing an N-terminal 6x-histidine-SUMO tag to facilitate downstream purification. Oligonucleotide primer pairs, each encoding for one half of the peptide sequences, were designed using NEBaseChanger V2, then incorporated into the plasmid using Q5 site-directed mutagenesis, as per the manufacturer’s instructions. Plasmid assembly was verified using Sanger sequencing (GENEWIZ) and then transformed into chemically competent Escherichia coli BL21(DE3) cells. Starter cultures (3 ml of LB media, 50 µg/ml kanamycin) were inoculated from freshly streaked agar plates or glycerol stocks and grown at 37°C with shaking at 225 r.p.m. overnight. Starter cultures were then diluted 1:500 in bulk cultures for Overnight Express Autoinduction media (Novagen), then grown overnight at 37 °C with shaking. Cells were then collected by centrifugation (4,500xg) at 4°C and washed twice with ice-cold 1x PBS. The resulting cell pellets were frozen at −20°C overnight, thawed to room temperature and then lysed using BugBuster protein extraction reagent (Millipore Sigma, 70584-3) supplemented with recombinant lysozyme (Millipore Sigma, 71110-3) and benzonase endonuclease (Millipore Sigma, E1014-25KU) for 20 minutes at room temperature with gentle rocking. The corresponding lysate was diluted with lysis buffer (1xPBS, 20 mM imidazole, 1x Halt protease inhibitor cocktail (Thermo Fisher Scientific, 78430)) and then centrifuged at 14,000xg for 30 minutes. The cleared supernatant was mixed end over end at 4°C for 30 minutes with HisPur Ni-NTA resin (Thermo Fisher Scientific, 88221) equilibrated with 20 mM imidazole in 1xPBS. Resin was centrifuged at 700xg for 2 minutes and then washed twice with 50 mM imidazole in 1xPBS, then twice more with 75 mM imidazole in 1x PBS. Protein was eluted with two consecutive washes using 500mM imidazole and desalted into PBS using Zeba spin desalting columns (Thermo Fisher Scientific, 89892). Expression and purity of purified proteins in both the soluble and insoluble fraction, as well as purified fractions, were assessed using SDS-PAGE (**Supplementary Figure 9**). Protein concentrations were quantified using a Qubit Protein Assay (Thermo Fisher Scientific, Q33211).

### Sandwich ELISAs

NCAM1 (2 ug/mL, Sino Biological 10673-H08H), NCAM1-FN3 (4 ug/mL) and β-catenin full length (2 ug/mL, Novus Biological NBP3-18198) were coated onto target protein were diluted in coating buffer (10 mM phosphate, pH 7.4), then coated onto 96-well plates at a volume of 50-100 uL per well at 4°C overnight with gentle rocking. Plates were washed once with Tris-buffered saline supplemented with 0.1% Tween-20 (v/v) (TBS-T) then blocked with 350µl of SuperBlock in PBS (Thermo Fisher Scientific, 37516) per the manufacturer’s instructions. moPPIt-derived peptides were serially diluted in biological triplicate using Superblock with 0.1% Tween-20, after which 100 uL of each solution was added to each well and incubated at room temperature for 1 hour. Plates were then washed five times using 350µl of TBS-T per well and then incubated with 100µl of SA-HRP (Thermo Fisher Scientific, N100, diluted 1:10,000 (NCAM1 assays) and 1:20,000 (β-catenin assays) in SuperBlock with 0.1% Tween-20) for 30 minutes at room temperature. Plates were again washed five times with 350µl of TBS-T and then incubated with 100µl per well of 3,3’-5,5’-tetramethylbenzidine substrate (1-Step Ultra TMB-ELISA; Thermo Fisher Scientific, 34029) for 15 minutes at room temperature with gentle rocking. Finally, the reaction was quenched with 100µl of 2M H2SO4, and absorbance at 450nm was immediately quantified using a Promega GloMax Discover plate reader.

### Primary monocyte isolation from PBMCs

Buffy coats from healthy donors were obtained from the Apheresis Suite of the Singapore Health Sciences Authority (HSA) in accordance with IRB guidelines. Buffy coats were kept on ice until processing. PBMCs were isolated by density gradient centrifugation using Ficoll-Paque PLUS (Cytiva, cat. no. 17144003). Briefly, buffy coats were diluted 1:1 with cold Hanks’ Balanced Salt Solution (HBSS; Gibco, cat. no. 14025092) and carefully layered over 10 mL of cold Ficoll-Paque PLUS in 50ml conical tubes. Samples were centrifuged at 400 x g for 20 min at 4°C with the brake set to the lowest setting. The interphase containing PBMCs was collected and washed twice with cold 1x phosphate-buffered saline (PBS) by centrifugation at 400 x g for 8 min. The resulting cell pellet was resuspended in 3 ml of TheraPEAK™ ACK lysing buffer (Lonza, cat. no. BP10-548E) for 5 min at room temperature to remove residual erythrocytes, followed by an additional PBS wash. The final PBMC pellet was resuspended in complete RPMI 1640 medium supplemented with 10% fetal bovine serum (FBS; 1st BASE, cat. no. CUL-1500-1L) and plated in T175 flasks. Monocytes were allowed to adhere for 24h at 37 °C in a humidified 5% CO_2_ incubator. Non-adherent cells were removed by gentle washing with warm 1x PBS, and adherent monocytes were detached by scraping and replated for subsequent assays.

### *In silico* generation of peptides targeting GM-CSFRα

Three classes of wild-type amino acid sequences were designed: (i) peptides targeting the pocket residues of the GM-CSF receptor (V50, E52, R54, R65, E66, S169, R170, K195, and R283) identified from the GM-CSF:GM-CSFRα crystal structure (PDB ID: 4RS1), (ii) motif-free peptides without predefined structural constraints, and (iii) peptides targeting the conserved WSxWS motif implicated in class I cytokine receptor activation (W287-S291).^65^ For each class, *in silico*-generated sequences were ranked using a compound score that favors high predicted motif- and affinity-scores, as well as solubility, while penalizing peptides with elevated predicted hemolytic activity. Peptides were ranked within each class according to this compound metric, and the top five candidates per class (15 total peptides) were commercially synthesized at 95% purity (Bio Basic). Peptides were received in lyophilised form, and resuspended according to the manufacturer’s guidelines. Peptides P4, MF4, MF5, and W4 did not solubilize well and were not included in the screen.

### Cell Morphology Assay

Primary human monocytes were isolated as described above and seeded into 96-well plates at approximately 8 x 10^3^ cells per well in complete RPMI 1640 culture medium. Monocytes were pre-incubated with 5 µM of the indicated peptides for 2 h, after which cells were stimulated with GM-CSF in the continued presence of 5 µM peptide. Culture medium containing GM-CSF and peptides was replenished every 24 h, and brightfield images of each well were acquired after 72 h of stimulation. Brightfield images were processed using the Cellpose segmentation algorithm to identify individual cells and define regions of interest (ROIs). For each condition, the mean projected cell area was calculated in Fiji from segmented ROIs and used as a quantitative surrogate of monocyte-to-macrophage differentiation, with larger, spread cell morphology corresponding to a more differentiated macrophage phenotype. Area scores were z-scaled by donor to account for donor-specific variability and provide a relative comparison to baseline sizes.

### Cloning of CAR constructs

A synthetic unique biologically and immunologically inert cell surface ligand was created by deleting the KTEL endoplasmic retention motif (DKTEL) and mutating the CXXS thioredoxin binding motif (C81S) of the anterior gradient protein 2 (AGR2) coding sequence (CDS),^45^ replacing the signal peptide (SP) sequence with that of the human kappa light chain CDS, and fusing this modified CDS to the transmembrane (TM) domain of platelet derived growth factor receptor (PDGFR). A chimeric antigen receptor (CAR) entry vector was created encoding a CD8A SP, Esp3I cloning site, Myc-tag, mutant long IgG4 hinge,^48^ CD28 TM domain, and CD28-CD3ζ signaling domain.^49^ The CAR gene was linked to a downstream GFP reporter gene via a 2A self-cleaving peptide sequence. For both constructs, gene expression was driven by an EF1A promoter, and the DNA was synthesized and subcloned in a lentiviral backbone by VectorBuilder (Chicago, IL). To assemble CARs featuring moPPIt-generated AGR2t-specific peptide binders as antigen binding domains, oligos for candidate peptide sequences were annealed and ligated via T4 DNA Ligase (New England Biolabs, Ipswich, MA) into the Esp3I-digested CAR lentiviral backbone.^54^ Assembled constructs were transformed into NEB OneShot Stbl3 Competent *Escherichia coli* bacteria (ThermoFisher Scientific, Waltham, MA) and plated onto LB agar plates supplemented with ampicillin for subsequent sequence verification of colonies and plasmid purification (Zymo Research, Irvine, CA). Construct sequences were verified by Sanger sequencing and whole plasmid sequencing (Eurofins Genomics, Louisville, KY).

### Lentivirus production and target cell line generation

HEK293FT cells were seeded at 3×10^6^ cells per 10 cm dish. The following morning, cells were treated with 25 mM chloroquine diphosphate. That afternoon, cells were transfected with DNA plasmids encoding lentiviral packaging proteins, lentiviral envelope proteins, and the construct of interest using polyethylenimine (PEI), as previously detailed.^66^ Lentivirus was harvested, concentrated, titrated in Jurkat cells, and frozen in single-use aliquots at −80°C until used. K562 cells were cultured in RPMI 1640 media supplemented with 10% fetal bovine serum (FBS), 1% Penicillin-Streptomycin, 2 mM GlutaMax, 10 mM HEPES, 1x non-essential amino acids (NEAA), and 1 mM sodium pyruvate (all from ThermoFisher Scientific). AGR2t-K562 target cells were generated by transducing K562 cells with AGR2t lentivirus, staining them with rabbit anti-human AGR2 antibody (Cell Signaling Technology, Danvers, MA) and goat anti-rabbit IgG AlexaFluor 647 (ThermoFisher Scientific), and sorting them to purity based on AGR2 surface expression using fluorescence activated cell sorting (FACS).

### Human regulatory T cell isolation, lentiviral transduction, and phenotyping

Chimeric antigen receptor modified regulatory T cells (CAR Tregs) were generated as previously detailed.^49,50^ De-identified healthy donor-derived human peripheral blood leukopaks were obtained from STEMCELL Technologies (Vancouver, Canada). CD4^+^ T cells were purified using the EasySep Human CD4^+^ T cell Isolation Kit (STEMCELL Technologies) following manufacturer’s instructions and stained for CD4 FITC, CD25 APC, and CD127 PE (all from Biolegend, San Diego, CA). CD4^+^CD25^high^CD127^low^ regulatory T cells (Tregs) and CD4^+^CD25^low^CD127^high^ effector T cells (Teff) were sorted by FACS using a BD FACS Aria II Cell Sorter (Beckton Dickinson, Franklin Lakes, NJ). Post-sort analyses confirmed population purity. Cells were counted with Trypan Blue using a TC20 Automated Cell Counter (BioRad, Hercules, CA) and seeded in 24-wells at 10^6^ cells per ml per 24-well with anti-CD3/CD28 beads (ThermoFisher Scientific) at a 1:1 bead to cell ratio and recombinant human IL-2 (ThermoFisher Scientific, Waltham, MA) in RPMI1640 medium supplemented with 10% FBS, 1% Penicillin-Streptomycin, 2 mM GlutaMax, 10 mM HEPES, 1x NEAA, and 1 mM sodium pyruvate. Tregs were cultured with 1,000 IU/mL IL-2, and total CD4^+^ T cells and CD4^+^ Teff cells with 100 IU/mL IL-2. Two days after activation with anti-CD3/CD28 and IL-2, T cells were counted and transduced with CAR lentivirus at a multiplicity of infection (MOI, lentivirus particles per cell) of 1 with IL-2. After adding CAR lentivirus, T cells were centrifuged at 1,000*g* at 32°C for 1h at 2.5×10^5^ cells per 1.5 ml Eppendorf tube. Following transduction, T cells were expanded in RPMI10 medium with IL-2. Transduction efficiency was assessed by flow cytometry based on Myc-tag and GFP expression at least 5 days post-transduction. To ascertain maintenance of Treg identity, CAR Tregs were intracellularly stained with FOXP3 APC (ThermoFisher Scientific) and HELIOS PE (Biolegend) using the eBioscience Foxp3/Transcription Factor Staining Buffer Set (ThermoFisher Scientific) per manufacturer’s instructions and analyzed using flow cytometry.

### Cardiomyocyte differentiation and culture

Human induced pluripotent stem cell-derived cardiomyocyte (iCM) differentiation was performed as previously described^67^ and according to the GiWi protocol^67^ to establish cardiac progenitor cells. In brief, 90% confluent 19-9-11 hiPSCs were singularized following incubation in Accutase for 6 minutes at 37°C, 5% CO_2_ and seeded onto Matrigel-coated 24-well plates at 600,000 cells/cm^2^ in mTeSR1 media with 5 µM Y-27632 on Day −2. Media was exchanged with mTeSR1 24h later (Day −1). On Day 0, cells were treated with 6-8 µM CHIR99021 (Selleck Chemicals, Houston, TX, USA) in RPMI 1640 (Thermo Fisher Scientific) supplemented with 2% B27 Minus Insulin (Thermo Fisher Scientific, Waltham, MA) (RPMI/B27-). Precisely 24 h later (Day 1), media was replaced with fresh RPMI/B27-. On Day 3, cells were treated with RPMI/B27-supplemented with IWP2 (Sigma-Aldrich). Precisely 48h later (Day 5), media was exchanged with fresh RPMI/B27-. By Day 6, cells were considered to be cardiac progenitor cells. On Day 7, media was replaced with RPMI 1640 supplemented with B27 with Insulin (Thermo Fisher Scientific) (RPMI/B27+). Media was subsequently replaced every other day. On Day 10, cells were purified via lactate purification^68^ for 48 h. Lactate purification media consisted of glucose-free DMEM (Thermo Fisher Scientific) supplemented with 3 mM Na-l-lactate (BeanTown Chemical, Hudson, NH, USA). On Day 12, media was replaced with RPMI/B27+ for 2 days of recovery from purification. On Day 14, purified hiPSC-CMs were prepared for expansion via dissociation using Accutase at 37°C, 5% CO_2_ for 40 minutes. Accutase was quenched 1:1 with RPMI/B27+ and centrifuged at 200*g* for 5 minutes at room temperature. iCMs were resuspended in RPMI/B27+ supplemented with 10% Defined FBS (Cytiva Life Sciences, Marlborough, MA, USA), and 5 µM Y-27632. Expansion of iCMs^69,70^ began 24 h after replating via replacement of media with RPMI/B27+ supplemented with 3 µM CHIR99021. Media was exchanged with RPMI/B27+ supplemented with 3 µM CHIR99021 every other day for cardiomyocyte expansion until iCMs grew to desired confluency.

### Cardiomyocyte lentiviral transduction

When ready for re-plating, iCMs were incubated in 1X TrypLE (Gibco) at 37°C, 5% CO_2_ for 40 minutes. TrypLE was quenched 1:1 with RPMI/B27+ and cells were centrifuged at 200*g* for 5 minutes at room temperature. iCMs were resuspended in RPMI/B27+ supplemented with 10% Defined FBS and 5 µM Y-27632 and plated onto Matrigel-coated 96-well plates at 20,000 cells/well. Media was exchanged with RPMI/B27+ every other day until lentiviral transduction. iCMs were transduced with AGR2t lentivirus at an MOI of 100 with polybrene. After adding AGR2t lentivirus, iCMs were centrifuged at 1,000*g* at 32°C for 1h.

### Chimeric antigen receptor T cell activation assay

CAR CD4^+^ T cells, CAR Tregs, and CAR Teff cells were co-cultured with parental or AGR2t-expressing target cells in RPMI10 medium. When using CAR Tregs, RPMI10 was supplemented with 1,000 IU/mL IL-2. CAR CD4^+^ T cells, CAR Tregs, and CAR Teff cells were seeded at 10^5^ cells per round-bottom 96-well with 10^5^ irradiated (3,000 rad) K562 or AGR2t-K562 cells or seeded at 10^5^ cells per flat-bottom 96-well containing 2×10^4^ iCMs or AGR2t-iCMs in 200 ml total volume per well. Two days later, co-cultures were harvested and surface expression of T cell activation markers CD25 and CD71 was assessed by flow cytometry by staining with CD4 PE-Cy7, CD25 APC, and CD71 PE (all from Biolegend). Untransduced (UT) and irrelevant CD19 CAR T cells were used as negative controls.

### Chimeric antigen receptor T cell-mediated cytotoxicity assay

CAR Teff cells were seeded at 10^5^ cells per flat-bottom 96-well containing 2×10^4^ iCMs or AGR2t-iCMs in 200 ml total volume per well. Two days later, 50 ml of supernatant was carefully collected from each well and analyzed using the CyQUANT Cytotoxicity Lactate Dehydrogenase Release Assay kit (Thermo Fisher Scientific) as per the manufacturer’s instructions to quantify target cell killing.^66^ UT Teff cells were included as negative controls. Undisturbed and lysed iCMs and AGR2t-iCMs seeded in adjacent wells were used to determine spontaneous and maximum LDH release, respectively.

### Statistical analysis and reproducibility

Statistical analyses for experimental methods were performed using GraphPad Prism v.10.6.1 (GraphPad Software, La Jolla, CA). Statistical measures are provided for each experiment in the Figure Legends. The authors were not blinded during the study.

## Author Contributions

T.C. and P.C. devised and developed model architectures and theoretical formulations. T.C. trained and benchmarked models. Z.Q. designed and performed all *in vitro* biophysical experiments. K.M. performed GM-CSF assays, with supervision from J.B. E.C.O, M.J.V., and L.M.R.F constructed and tested AGR2t-targeting CARs, which were validated in cardiomyocytes generated by S.E.S. and Y.M. Y.Z. advised on model design and theoretical framework, and helped to process data for training. Z.Q., T.C., and P.C. wrote the manuscript, with input from all authors. L.M.R.F devised and supervised the entire CAR Treg work. P.C. designed, supervised, and directed the study, and reviewed and finalized the manuscript.

## Data and Materials Availability

The datasets and codebase to train BindEvaluator and construct moPPIt is freely accessible at https://huggingface.co/ChatterjeeLab/moPPIt, alongside an easy-to-use Colab notebook for inference (https://colab.research.google.com/drive/16n8PIwKwAiG-oDLm171BWvv-lQH0dHMg?usp=sharing). All raw And processed experimental data can be found at our Zenodo repository: https://zenodo.org/records/18008663.

## Acknowledgements

The authors would like to thank Sophia Tang for advice on theoretical formulations and Lauren Hong for logo development.

## Declarations

The research was supported by The Hartwell Foundation and NIH grants 3U54CA231630-01A1S4 and 1R21CA278468-01 to the lab of P.C, NIH grants 1R01HL175050 and UL1TR001450, and NSF grant #OIA-2242812 to the lab of Y.M., NIH grants U24DK104162-07, UL1TR001450, and 1R01HL175050 to the lab of L.M.R.F. The work was also supported in part by the Flow Cytometry and Cell Sorting Shared Resource, Hollings Cancer Center, Medical University of South Carolina (P30 CA138313). J.B. acknowledges funding from the National Medical Research Council (OFIRG24jan-0083/ MOH-001622 and OFLCG22may-0011/ MOH-001327-05), Ministry of Education Singapore (MOE-T2EP30125-0010), Agency for Science and Technology (Industry Alignment Fund Prepositioning grant, H24J4a0019) and Duke NUS (Duke-NUS-GCR/2024/0034).

