## Supplementary Information for "moPPIt: *De Novo* Generation of Motif-Specific and Functionally Active Peptide Binders via Discrete Flow Matching"

**Supplementary Figures**

1. Validation loss curves for BindEvaluator training and fine-tuning.
2. PeptiDerive relative interface scores for existing and designed peptide-protein complexes (Set 1).
3. PeptiDerive relative interface scores for existing and designed peptide-protein complexes (Set 2).
4. Structural visualization and PeptiDerive relative interaction scores for designed peptides targeting structured motifs (Set 1).
5. Structural visualization and PeptiDerive relative interaction scores for designed peptides targeting structured motifs (Set 2).
6. Structural visualization and PeptiDerive relative interaction scores for designed peptides targeting structured motifs (Set 3).
7. Structural visualization and PeptiDerive relative interaction scores for designed peptides targeting intrinsically disordered proteins.
8. Cell line characterization, controls, and gating strategy used for CAR T cell assays.

**Supplementary Tables**

1. Peptide sequences (with associated metrics) used for *in silico* validation.
2. Peptide sequences (with associated metrics) used for experimental validation.
3. Target protein sequences used in this study.

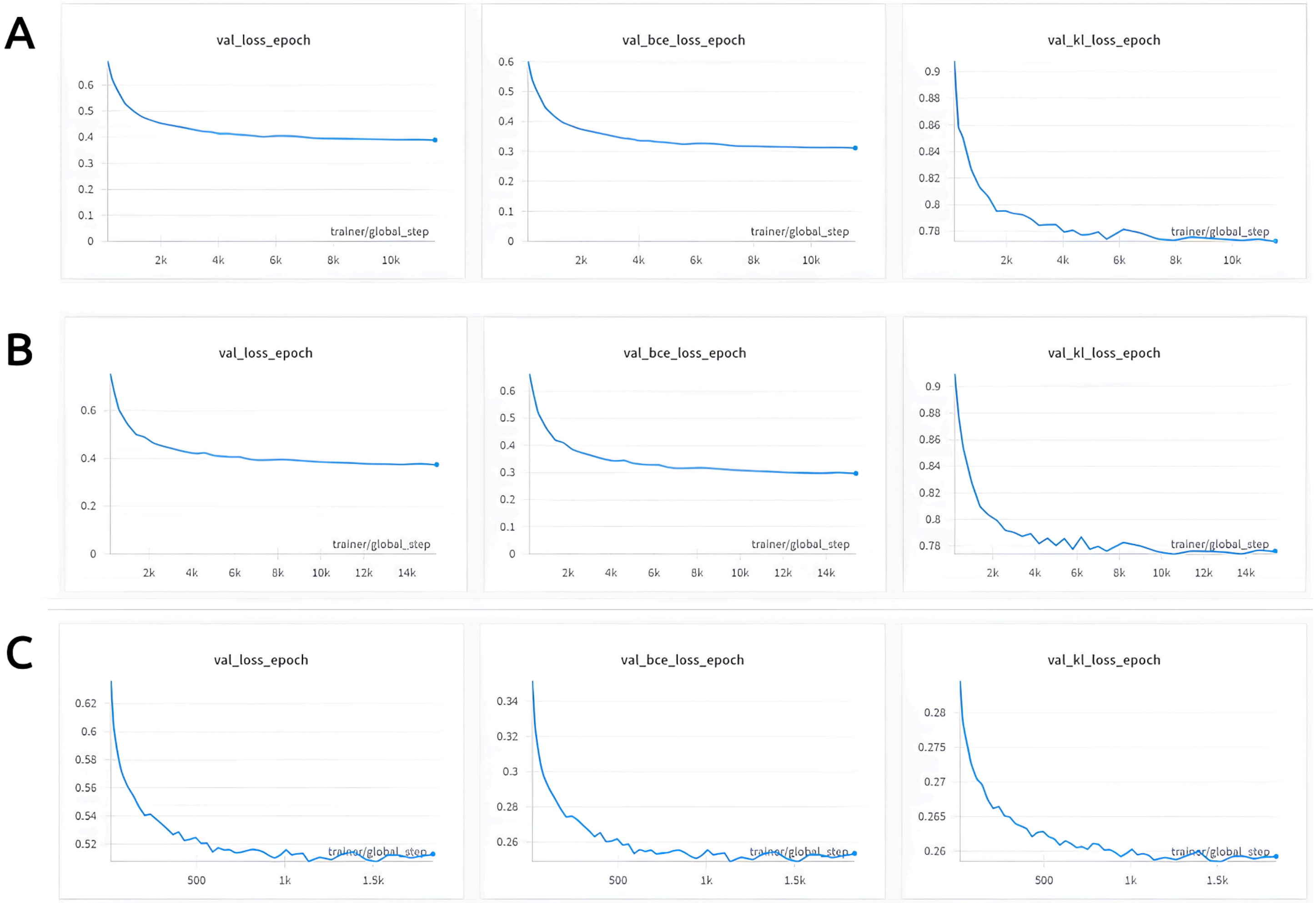


**Supplementary Figure 1: Validation loss curves for BindEvaluator training and fine-tuning. (A)** Validation loss, binary cross-entropy (BCE) loss, and Kullback-Leibler (KL) divergence loss curves during training of BindEvaluator on the PPI dataset without dilated CNN modules. **(B)** Loss curves for training with dilated CNN modules, showing similar trends to (A) but with noticeable reductions in losses during the final epochs. **(C)** Loss curves during fine-tuning of BindEvaluator with dilated CNN modules on peptide-protein binding data, illustrating further decreases in loss metrics, particularly in KL divergence.

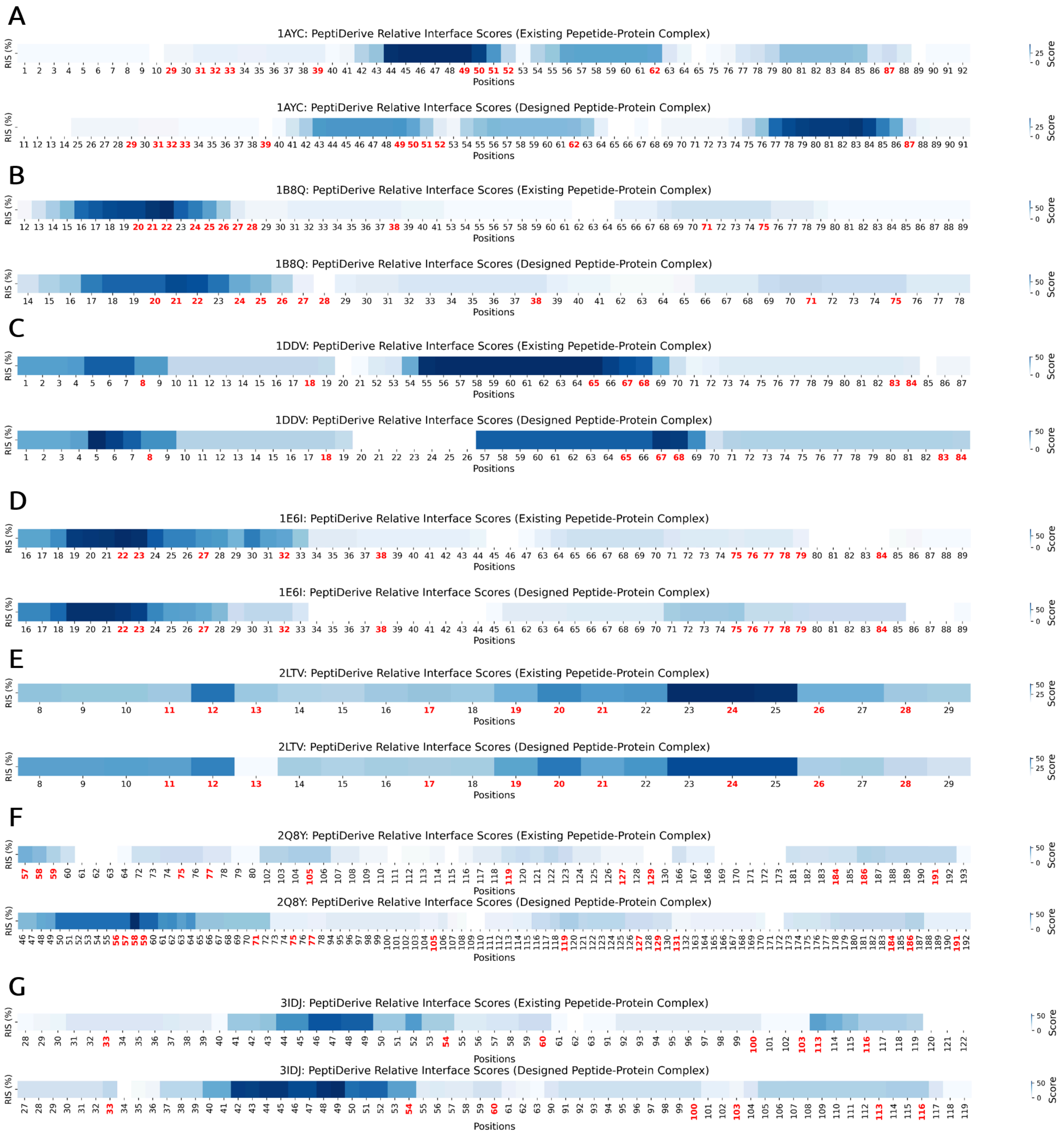


**Supplementary Figure 2: PeptiDerive relative interface scores for existing and designed peptide-protein complexes (Set 1).** Heatmaps of PeptiDerive relative interface scores (RIS) are shown for 7 peptide-protein complexes among 15 structured complexes with known binders that were tested: **(A)** 1AYC, **(B)** 1B8Q, **(C)** 1DDV, **(D)** 1E6I, **(E)** 2LTV, **(F)** 2Q8Y, **(G)** 3IDJ. The first heatmap for each protein shows the RIS of the existing peptide-protein complex, while the second heatmap shows the scores for the designed peptide-protein complex. For each heatmap, the x-axis indicates the residue positions, with highlighted positions in red representing the target binding amino acid positions that were input into moPPIt. High RIS at these positions indicate strong binding potential.

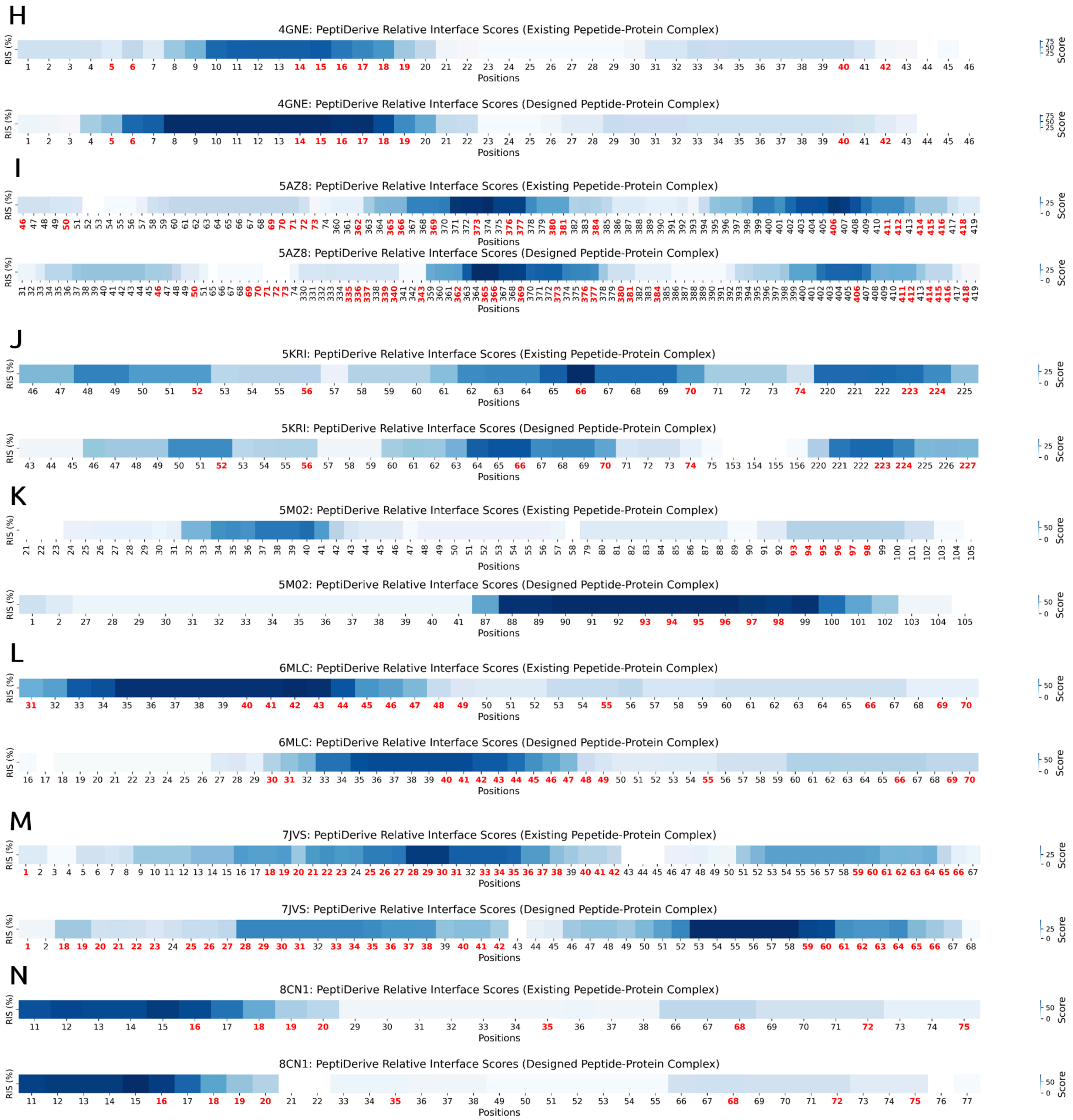


**Supplementary Figure 3: PeptiDerive relative interface scores for existing and designed peptide-protein complexes (Set 2).** Heatmaps of PeptiDerive relative interface scores (RIS) are shown for 7 peptide-protein complexes among 15 structured complexes with known binders that were tested: **(H)** 4GNE, **(I)** 5AZ8, **(J)** 5KRI, **(K)** 5M02, **(L)** 6MLC, **(M)** 7JVS, **(N)** 8CN1. The first heatmap for each protein shows the RIS of the existing peptide-protein complex, while the second heatmap shows the scores for the designed peptide-protein complex. For each heatmap, the x-axis indicates the residue positions, with highlighted positions in red representing the target binding amino acid positions that were input into moPPIt. High RIS at these positions indicate strong binding potential.

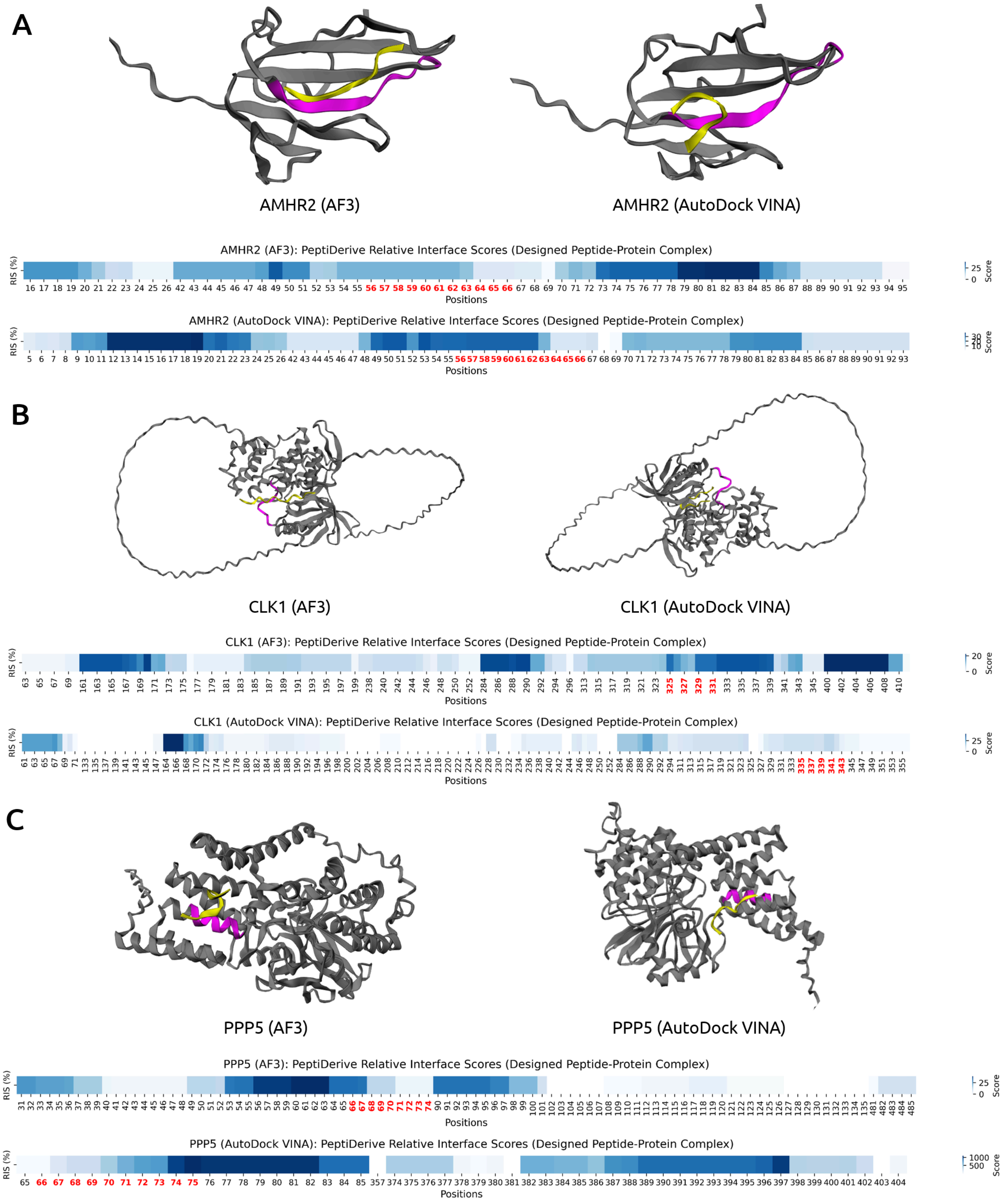


**Supplementary Figure 4: Structural visualization and PeptiDerive relative interaction scores for designed peptides targeting structured motifs (Set 1).** The peptide-protein complex structures are visualized for three proteins without known binders: **(A)** AMHR2, **(B)** CLK1, **(C)** PPP5 using AlphaFold3 and AutoDock VINA. The target proteins are depicted in grey, the designed peptide binders are shown in yellow, and the binding residues specified by the moPPIt-v3 algorithm are highlighted in magenta. Below each structure, the relative interaction scores (RIS) computed by PeptiDerive are shown, with high scores indicating strong binding potential. Positions highlighted in red were input into moPPIt-v3 as the desired target amino acids.

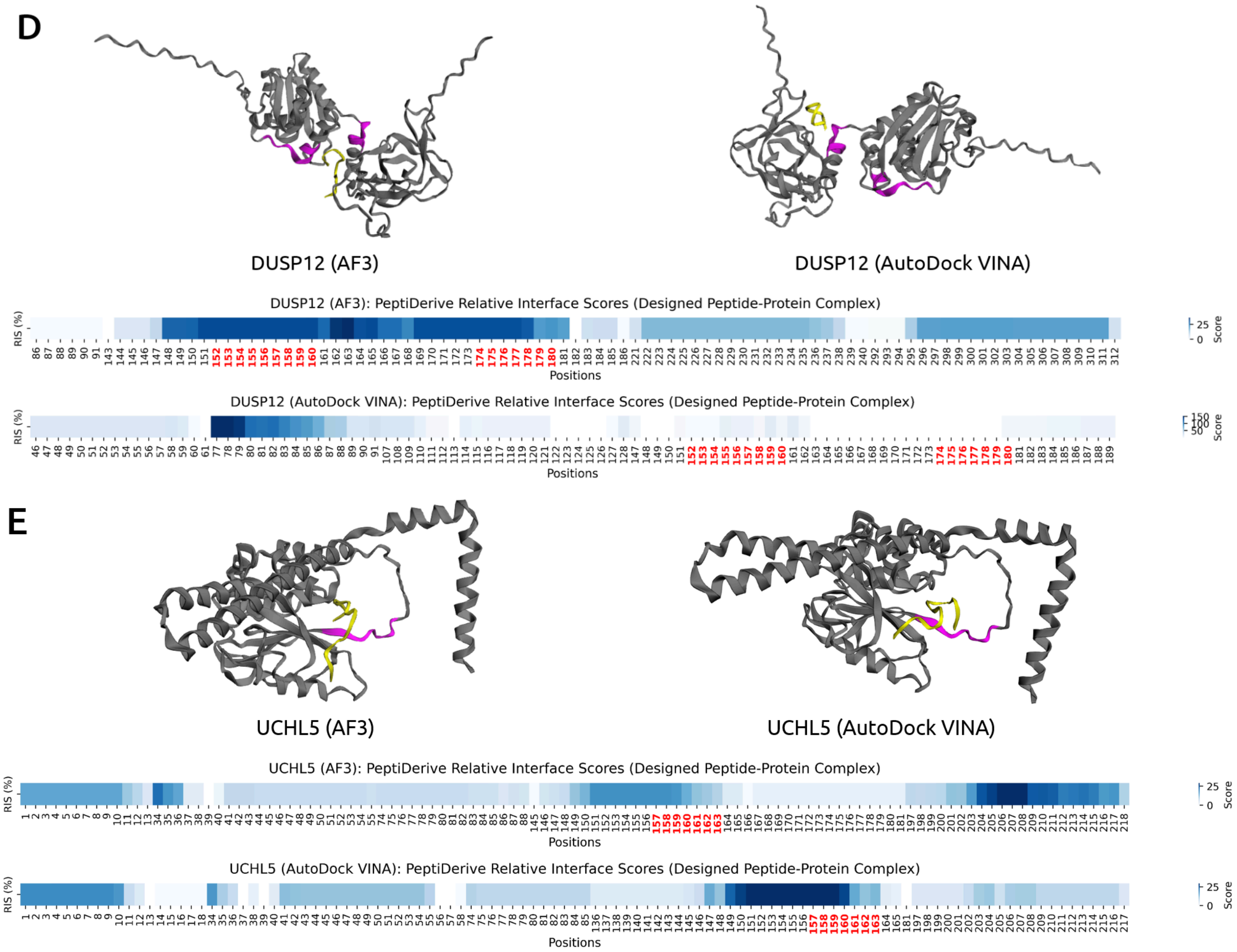


**Supplementary Figure 5: Structural visualization and PeptiDerive relative interaction scores for designed peptides targeting structured motifs (Set 2).** The peptide-protein complex structures are visualized for two proteins without known binders: **(D)** DUSP12, **(E)** UCHL5 using AlphaFold3 and AutoDock VINA. The target proteins are depicted in grey, the designed peptide binders are shown in yellow, and the binding residues specified by the moPPIt-v3 algorithm are highlighted in magenta. Below each structure, the relative interaction scores (RIS) computed by PeptiDerive are shown, with high scores indicating strong binding potential. Positions highlighted in red were input into moPPIt-v3 as the desired target amino acids.

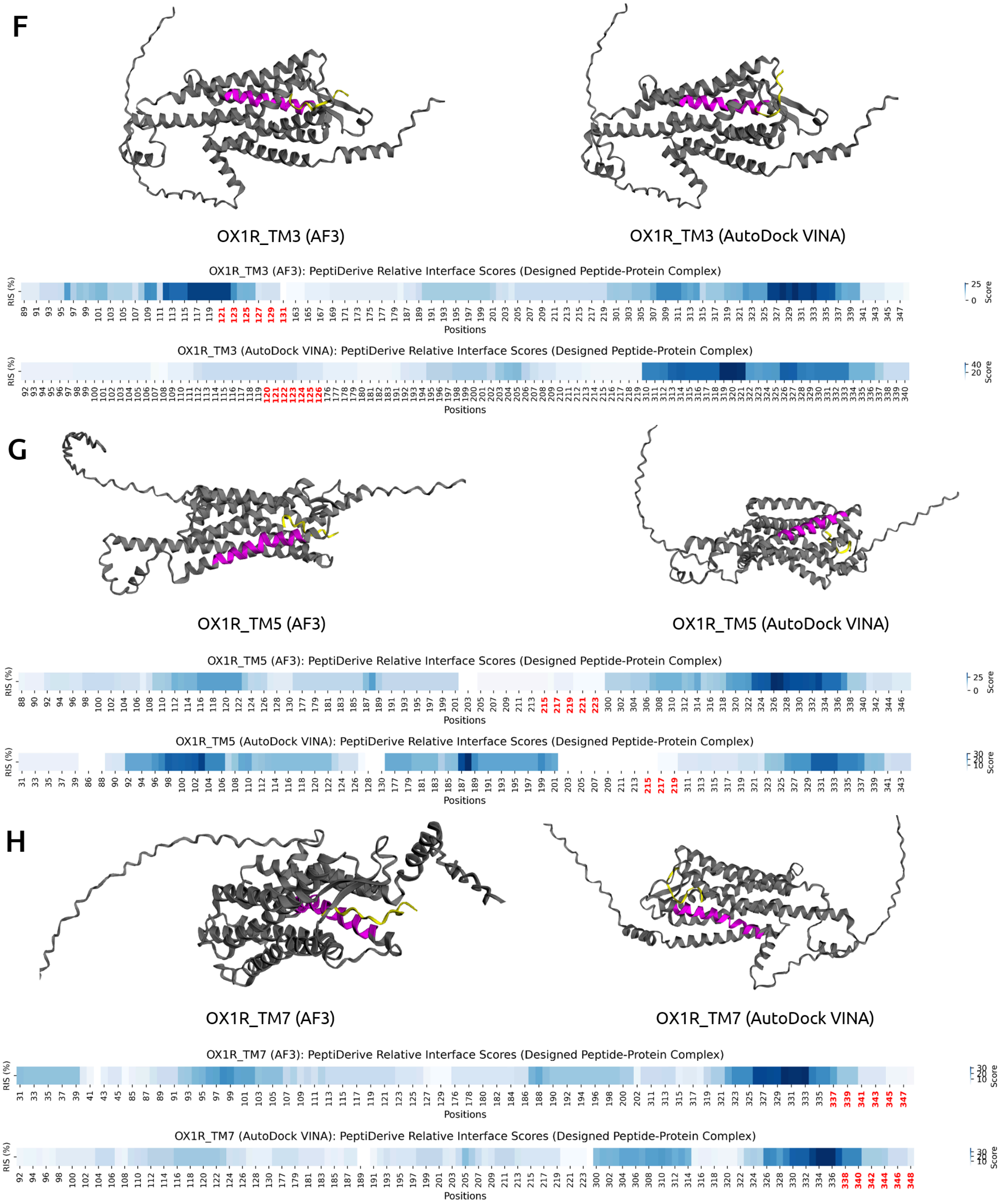


**Supplementary Figure 6: Structural visualization and PeptiDerive relative interaction scores for designed peptides targeting structured motifs (Set 3).** The peptide-complex structures are visualized for three different domains on OX1R: **(F)** Transmembrane (Name=3), **(G)** Transmembrane (Name=5), **(H)** Transmembrane (Name=7) using AlphaFold3 and AutoDock VINA. The target proteins are depicted in grey, the designed peptide binders are shown in yellow, and the binding residues specified by the moPPIt-v3 algorithm are highlighted in magenta. Below each structure, the relative interaction scores (RIS) computed by PeptiDerive are shown, with high scores indicating strong binding potential. Positions highlighted in red were input into moPPIt-v3 as the desired target amino acids.

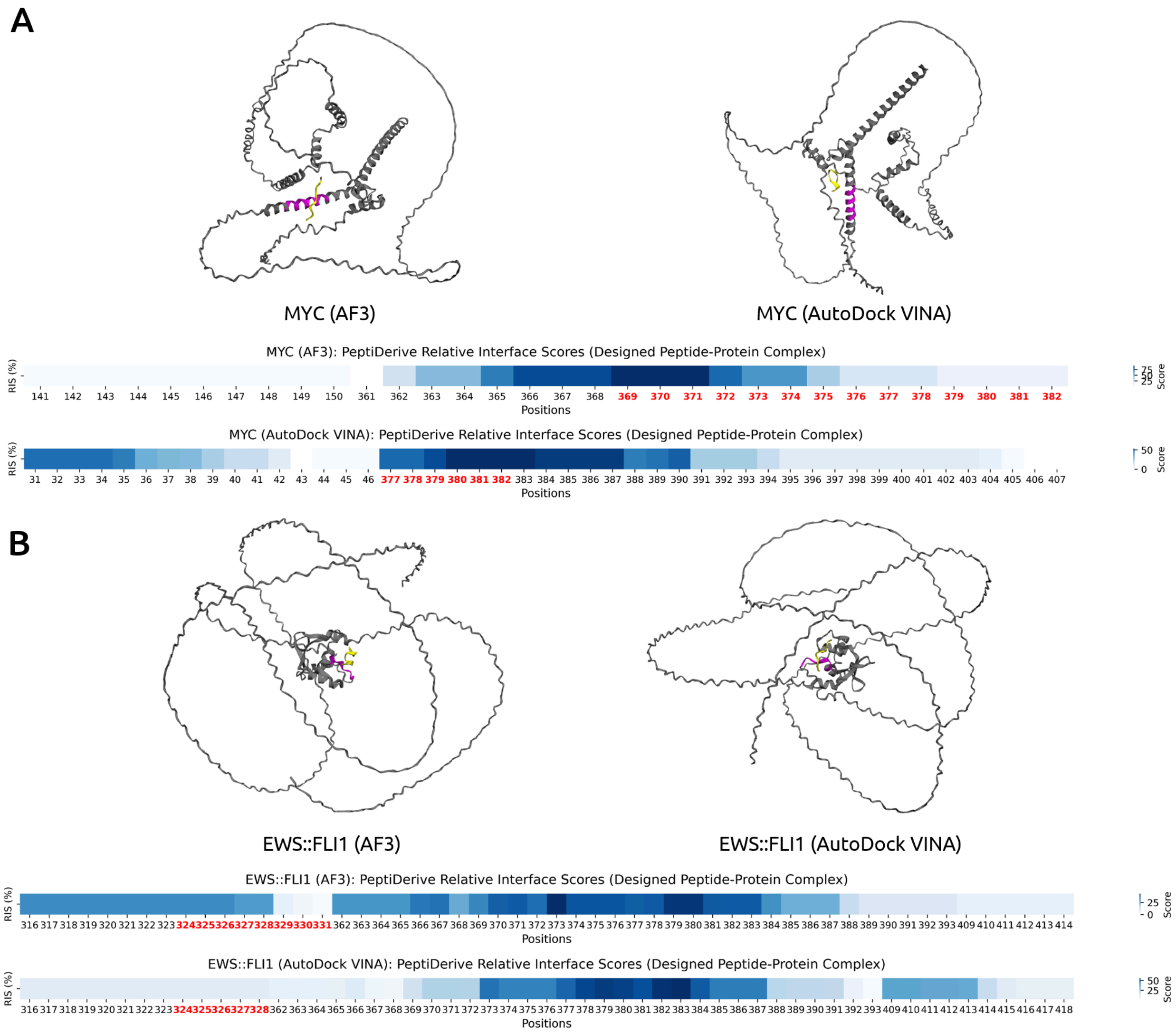


**Supplementary Figure 7: Structural visualization and PeptiDerive relative interaction scores for designed peptides targeting intrinsically disordered proteins.** The peptide-complex structures are visualized for two intrinsically disordered proteins: **(A)** MYC, **(B)** EWS::FLI1 using AlphaFold3 and AutoDock VINA. The target proteins are depicted in grey, the designed peptide binders are shown in yellow, and the binding residues specified by the moPPIt-v3 algorithm are highlighted in magenta. Below each structure, the relative interaction scores (RIS) computed by PeptiDerive are shown, with high scores indicating strong binding potential. Positions highlighted in red were input into moPPIt-v3 as the desired target amino acids.

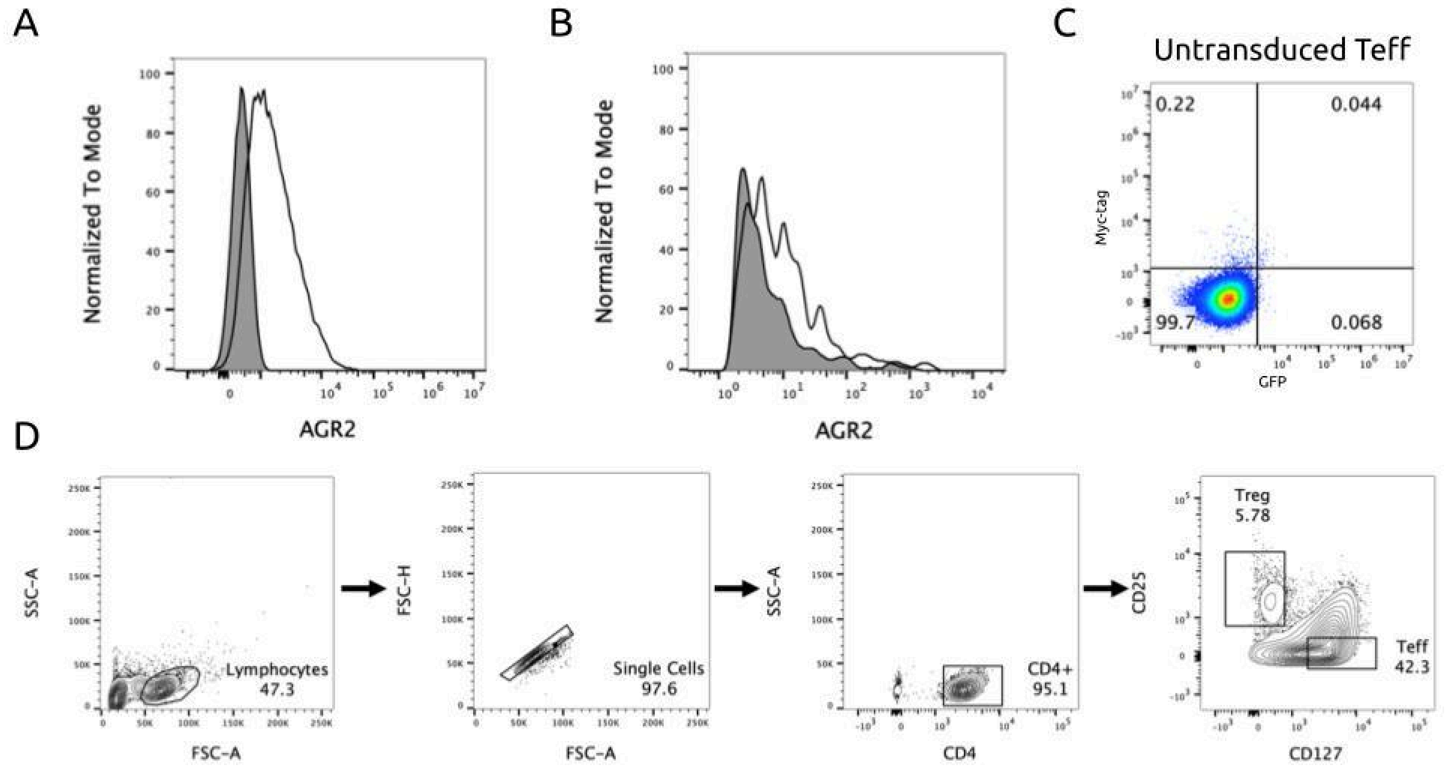


**Supplementary Figure 8. Cell line characterization, controls, and gating strategy used for CAR T cell assays. (A)** Surface expression of AGR2t in K562 cells. Shaded histogram represents untransduced K562 cells and line histogram represents AGR2t lentivirus transduced K562 cells. (**B)** Surface expression of AGR2t in human induced pluripotent stem cell-derived cardiomyocytes (iCM). Shaded histogram represents untransduced iCMs and line histogram represents AGR2t lentivirus transduced iCMs. **(C)** Effector T-cell (Teff) controls demonstrating no AGR2t_moPPIt_M2 CAR surface expression (Myc-tag) and GFP reporter expression prior to lentiviral transduction. **(D)** Gating strategy to sort $CD4^{+}CD25^{+}CD127^{-}$ regulatory T cells (Treg) and $CD4^{+}CD25^{-}CD127^{+}$ effector T cells (Teff) from human $CD4^{+}$ T cells using fluorescence activated cell sorting (FACS).

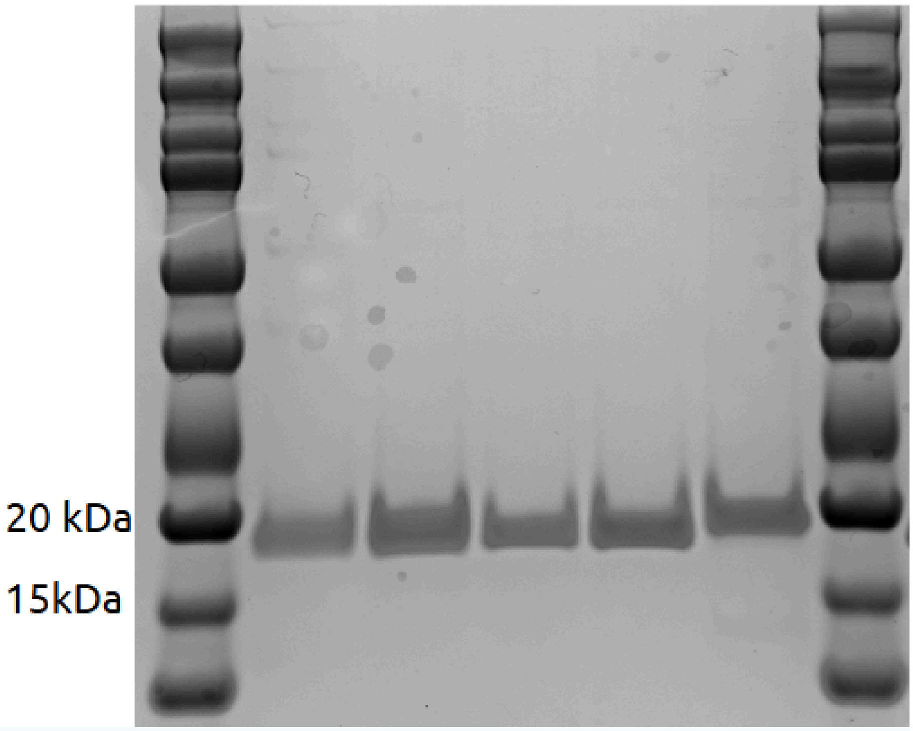


**Supplementary Figure 9. Representative coomassie-stained sodium dodecyl sulfate polyacrylamide gel electrophoresis (SDS-PAGE) of peptide constructs expressed as C-terminal SUMO tag fusions.** While constructs have a predicted molecular weight of 14.2 kDa based on amino acid sequence, the addition of a SUMO-tag fusion results in a higher apparent molecular weight of around 20 kDa.

**Supplementary Table 1:** Peptide sequences (with associated metrics) used for *in silico* validation.

| PDB ID | ipTM score (existing binder) | ipTM score (designed binder) | VINA score (existing binder) kcal/mol | VINA score (designed binder) kcal/mol | Designed Binder |
|---|---|---|---|---|---|
| 1AYC | 0.52 | 0.64 | -6.1 | -4.9 | YAYRYICYYCD |
| 1B8Q | 0.72 | 0.72 | -5.1 | -5.6 | IVDWVCF |
| 1DDV | 0.56 | 0.89 | -6.1 | -6.7 | RCVRWC |
| 1E6I | 0.58 | 0.66 | -7.4 | -7 | GRWRC |
| 2LTV | 0.56 | 0.6 | -3.4 | -4.6 | PTVEECSYWYHE |
| 2Q8Y | 0.52 | 0.69 | -7.3 | -5.2 | WLSWCHVYC |
| 3IDJ | 0.66 | 0.69 | -6.3 | -6.5 | IRRVRAP |
| 4GNE | 0.88 | 0.83 | -5.4 | -5.3 | ARRVRWS |
| 5AZ8 | 0.71 | 0.8 | -7.2 | -6.9 | LRWEVYLVREV |
| 5KRI | 0.85 | 0.84 | -3.8 | -3.6 | FAGMIVVNCIMR |
| 5M02 | 0.55 | 0.6 | -6.1 | -4.1 | PEVRWEVRD |
| 6MLC | 0.73 | 0.8 | -5.5 | -6.6 | GRWYCW |
| 7LUL | 0.94 | 0.89 | -7.2 | -6.8 | WEVTIWV |
| 7JVS | 0.43 | 0.54 | -5.5 | -8.1 | CVGIICEIICP |
| 8CN1 | 0.94 | 0.93 | -6.2 | -6 | SAEV |

**Supplementary Table 2:** Peptide sequences (with associated metrics) used for experimental validation.

| Peptide name | Peptide sequence | Target protein | Target motif | Motif Score | Affinity Score |
|---|---|---|---|---|---|
| NCAM1_moPPI t_F1 | PIMPNPPECDYA | NCAM1 | FN3 domain 2 | 0.30 | 5.10 |
| NCAM1_moPPIt_F2 | KDWNTIEEPT | NCAM1 | FN3 domain 2 | 0.31 | 4.94 |
| NCAM1_moPPIt_F3 | TEDYITCPPE | NCAM1 | FN3 domain 2 | 0.34 | 6.42 |
| NCAM1_moPPIt_F4 | DEQSMYIPPHDH | NCAM1 | FN3 domain 2 | 0.31 | 6.34 |
| βcat_moPPIt_1 | DQFEDEIEIF | β-catenin | N-terminal IDR | 0.46 | 0.83 |
| βcat_moPPIt_2 | FEEELFNIPIYD | β-catenin | N-terminal IDR | 0.55 | 0.80 |
| βcat_moPPIt_3 | ITFEFPIIIP | β-catenin | N-terminal IDR | 0.45 | 0.83 |
| βcat_moPPIt_4 | IIFPLFPFEE | β-catenin | N-terminal IDR | 0.52 | 0.81 |
| βcat_moPPIt_5 | YGTYGGFMVW | β-catenin | N-terminal IDR | 0.62 | 0.77 |
| βcat_moPPIt_6 | GMGMGTTGYQ | β-catenin | N-terminal IDR | 0.66 | 0.65 |
| βcat_moPPIt_7 | GYGTTFVTYCIF | β-catenin | N-terminal IDR | 0.52 | 0.78 |
| βcat_moPPIt_8 | EFDMGDMLGQ | β-catenin | N-terminal IDR | 0.72 | 0.63 |
| βcat_moPPIt_9 | FGGYDPFGGNIE | β-catenin | N-terminal IDR | 0.70 | 0.62 |
| βcat_moPPIt_10 | DTGIYGGFCYNC | β-catenin | N-terminal IDR | 0.70 | 0.76 |
| GMCSF_moPPIt_P1 | VMGFKIRVTHK | GM-CSF | Pocket | 0.69 | 7.86 |
| GMCSF_moPPIt_P2 | TVFLHRWVTKVTKLIRKVRD | GM-CSF | Pocket | 0.72 | 7.47 |
| GMCSF_moPPIt_P3 | TMCKVIKGIKL | GM-CSF | Pocket | 0.64 | 6.85 |
| GMCSF_moPPIt_P4 | VYMEVVKPQTYMCAWVLA | GM-CSF | Pocket | 0.57 | 7.36 |
| GMCSF_moPPIt_P5 | KAIGKKTRLYMKVIV | GM-CSF | Pocket | 0.68 | 6.84 |
| GMCSF_moPPIt_WS1 | PVEQCTVVTFVVERK | GM-CSF | WSxWS | 0.80 | 8.39 |
| GMCSF_moPPIt_WS2 | ISYAVVTKKK | GM-CSF | WSxWS | 0.73 | 7.92 |
| GMCSF_moPPIt_WS3 | PTERVAKIVCVKLKFG | GM-CSF | WSxWS | 0.79 | 7.19 |
| GMCSF_moPPIt_WS4 | GTGCAAAEIRVAVMQFF | GM-CSF | WSxWS | 0.77 | 7.44 |
| GMCSF_moPPIt_WS5 | DTVVREAVVKYCVQRV | GM-CSF | WSxWS | 0.86 | 7.95 |
| GMCSF_moPPIt_MF1 | ITTATRVEVKCK | GM-CSF | None | NA | 8.44 |
| GMCSF_moPPIt_MF2 | IKFTWEVTMTKVRAW | GM-CSF | None | NA | 8.28 |
| GMCSF_moPPIt_MF3 | WMTTPDIVVGSPEVRTMRK | GM-CSF | None | NA | 8.28 |
| GMCSF_moPPIt_MF4 | ARNTWVVTSGCMGV | GM-CSF | None | NA | 7.98 |
| GMCSF_moPPIt_MF5 | ITKTFACVGVDVGV | GM-CSF | None | NA | 8.06 |
| AGR2t_moPPIt_M2 | LVPLVVEVWVPD | AGR2t | None | 0.31 | 7.48 |
| AGR2t_moPPIt_M3 | AELVRFPCEVEC | AGR2t | None | 0.29 | 8.45 |
| AGR2t_moPPIt_M4 | DRVEVEDSPEIV | AGR2t | None | 0.27 | 9.08 |
| AGR2t_moPPItt_M5 | RKQSETIENSYP | AGR2t | None | 0.26 | 5.12 |

**Supplementary Table 3:** Target protein sequences used in this study.

**NCAM1 protein sequence**

MLQTKDLIWTLFFLGTAVSLQVDIVPSQGEISVGESKFFLCQVAGDAKDKDISWFSPNGEKLTPNQQRISVVWN
DDSSSTLTIYNANIDDAGIYKCVVTGEDGSESEATVNVKIFQKLMFKNAPTPQEFREGEDAVIVCDVVSSLPPTII
WKHKGRDVILKKDVRFIVLSNNYLQIRGIKKTDEGTYRCEGRILARGEINFKDIQVIVNVPPTIQARQNIVNATANL
GQSVTLVCDAEGFPEPTMSWTKDGEQIEQEEDDEKYIFSDDSSQLTIKKVDKNDEAEYICIAENKAGEQDATIHL
KVFAKPKITYVENQTAMELEEQVTLTCEASGDPIPSITWRTSTRNISSEEKASWTRPEKQETLDGHMVVRSHAR
VSSLTLKSIQYTDAGEYICTASNTIGQDSQSMYLEVQYAPKLQGPVAVYTWEGNQVNITCEVFAYPSATISWFRD
GQLLPSSNYSNIKIYNTPSASYLEVTPDSENDFGNYNCTAVNRIGQESLEFILVQADTPSSPSIDQVEPYSSTAQV
QFDEPEATGGVPILKYKAEWRAVGEEVWHSKWYDAKEASMEGIVTIVGLKPETTYAVRLAALNGKGLGEISAAS
EFKTQPVQGEPSAPKLEGQMGEDGNSIKVNLIKQDDGGSPIRHYLVRYRALSSEWKPEIRLPSGSDHVMLKSL
DWNAEYEVYVVAENQQGKSKAAHFVFRTSAQPTAIPANGSPTSGLSTGAIVGILIVIFVLLLVVVDITCYFLNKCG
LFMCIAVNLCGKAGPGAKGKDMEEGKAAFSKDESKEPIVEVRTEEERTPNHDGGKHTEPNETTPLTEPEKGPV
EAKPECQETETKPAPAEVKTVPNDATQTKENESKA

Immunoglobulin-like domain 2

Fibronectin type-3 domain 2

**β-catenin protein sequence**

MATQADLMELDMAMEPDRKAAVSHWQQQSYLDSGIHSGATTTAPSLSGKGNPEEEDVDTSQVLYEWEQGFS
QSFTQEQVADIDGQYAMTRAQRVRAAMFPETLDEGMQIPSTQFDAAHPTNVQRLAEPSQMLKHAVVNLINYQ
DDAELATRAIPELTKLLNDEDQVVVNKAAVMVHQLSKKEASRHAIMRSPQMVSAIVRTMQNTNDVETARCTAGT
LHNLSHHREGLLAIFKSGGIPALVKMLGSPVDSVLFYAITTLHNLLLHQEGAKMAVRLAGGLQKMVALLNKTNVK
FLAITTDCLQILAYGNQESKLIILASGGPQALVNIMRTYTYEKLLWTTSRVLKVLSVCSSNKPAIVEAGGMQALGL
HLTDPSQRLVQNCLWTLRNLSDAATKQEGMEGLLGTLVQLLGSDDINVVTCAAGILSNLTCNNYKNKMMVCQV
GGIEALVRTVLRAGDREDITEPAICALRHLTSRHQEAEMAQNAVRLHYGLPVVVKLLHPPSHWPLIKATVGLIRNL
ALCPANHAPLREQGAIPRLVQLLVRAHQDTQRRTSMGGTQQQFVEGVRMEEIVEGCTGALHILARDVHNRIVIR
GLNTIPLFVQLLYSPIENIQRVAAGVLCELAQDKEAAEAIEAEGATAPLTELLHSRNEGVATYAAAVLFRMSEDKP
QDYKKRLSVELTSSLFRTEPMAWNETADLGLDIGAQGEPLGYRQDDPSYRSFHSGGYGQDALGMDPMMEHE
MGGHHPGADYPVDGLPDLGHAQDLMDGLPPGDSNQLAWFDTDL

N-terminal disordered domain

**GM-CSF protein sequence**

MWLQNLLLLGAVVCSISAPTRLPSPVTRPWQHVDAIKEALSLLNNSNDTAAVMNETVDVVCKMFDPQEPTCVQ
TRLNLYKQGLRGSLTRLKSPLTLLAKHYEQHCPLTEETSCETQSITFKSFKDSLNKFLFTIPFDCWGPVKK

**AGR2t protein sequence**

METDTLLLWVLLLWVPGSTGDRDTTVKPGAKKDTKDSRPKLPQTLSRGWGDQLIWTQTYEEALYKSKTSNKPL
MIIHHLDE**SPHS**QALKKVFAENKEIQKLAEQFVLLNLVYETTDKHLSPDGQYVPRIMFVDPSLTVRADITGRYSNR
LYAYEPADTALLLDNMKKALKLLNAVGQDTQEVIVVPHSLPFKVVVISAILALVVLTIISLIILIMLWQKKPR

IgK signal peptide

AGR2 C18S deltaKTEL (**bolded** section represents the mutated thioredoxin-like domain SPHS is a mutant inactive form of the CPHS domain in wild-type AGR2 obtained by mutating the cysteine C into a serine (S)). . The sequence above is also missing the C-terminal endoplasmic reticulum (ER) retention KTEL amino acid sequence.

PDGFR TM domain (platelet derived growth factor receptor transmembrane)
