## Supplementary material for "moPPIt: *De Novo* Generation of Motif-Specific and Functionally Active Peptide Binders via Discrete Flow Matching": Figure 1

A

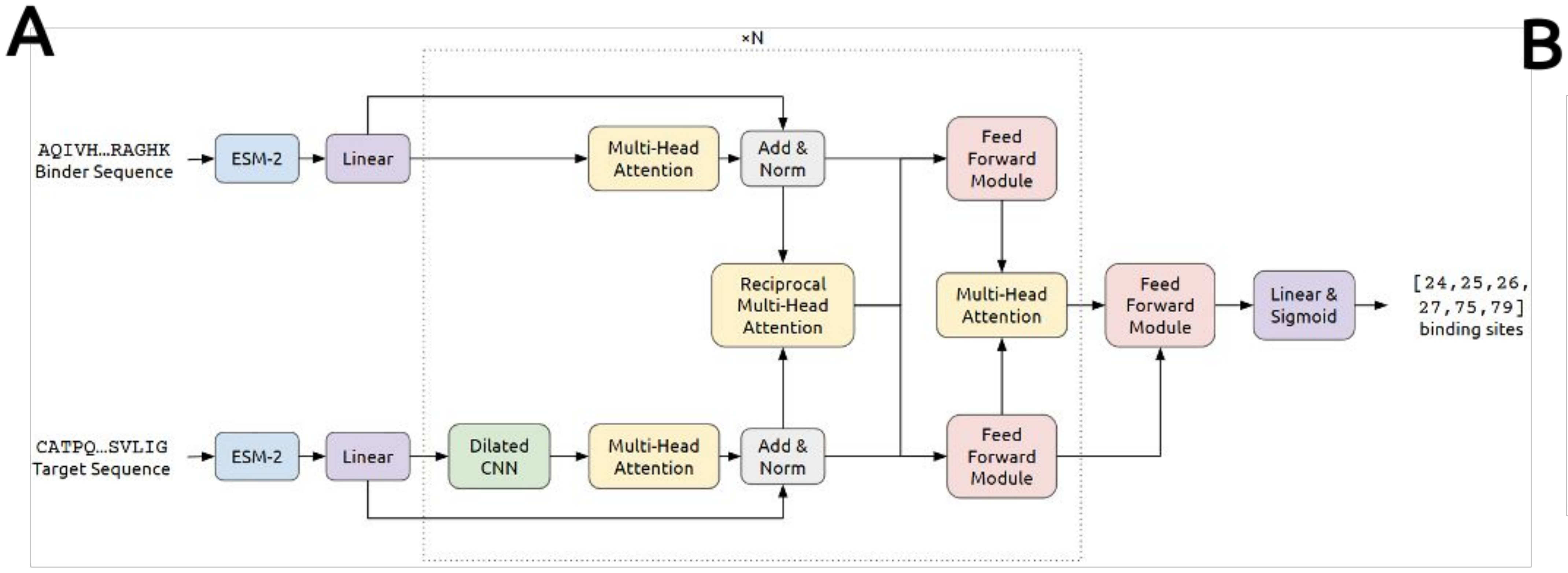


B

| Test Metric | Train w/o CNN | Train w/ CNN | Fine-tune w/ CNN |
|---|---|---|---|
| Loss | 0.388 | 0.373 | 0.514 |
| BCE Loss | 0.311 | 0.295 | 0.580 |
| KL Loss | 0.773 | 0.776 | 0.254 |
| Accuracy | 0.83 | 0.84 | 0.91 |
| AUC | 0.93 | 0.94 | 0.97 |
| F1 Score | 0.65 | 0.66 | 0.58 |
| MCC | 0.59 | 0.61 | 0.59 |

C D E F

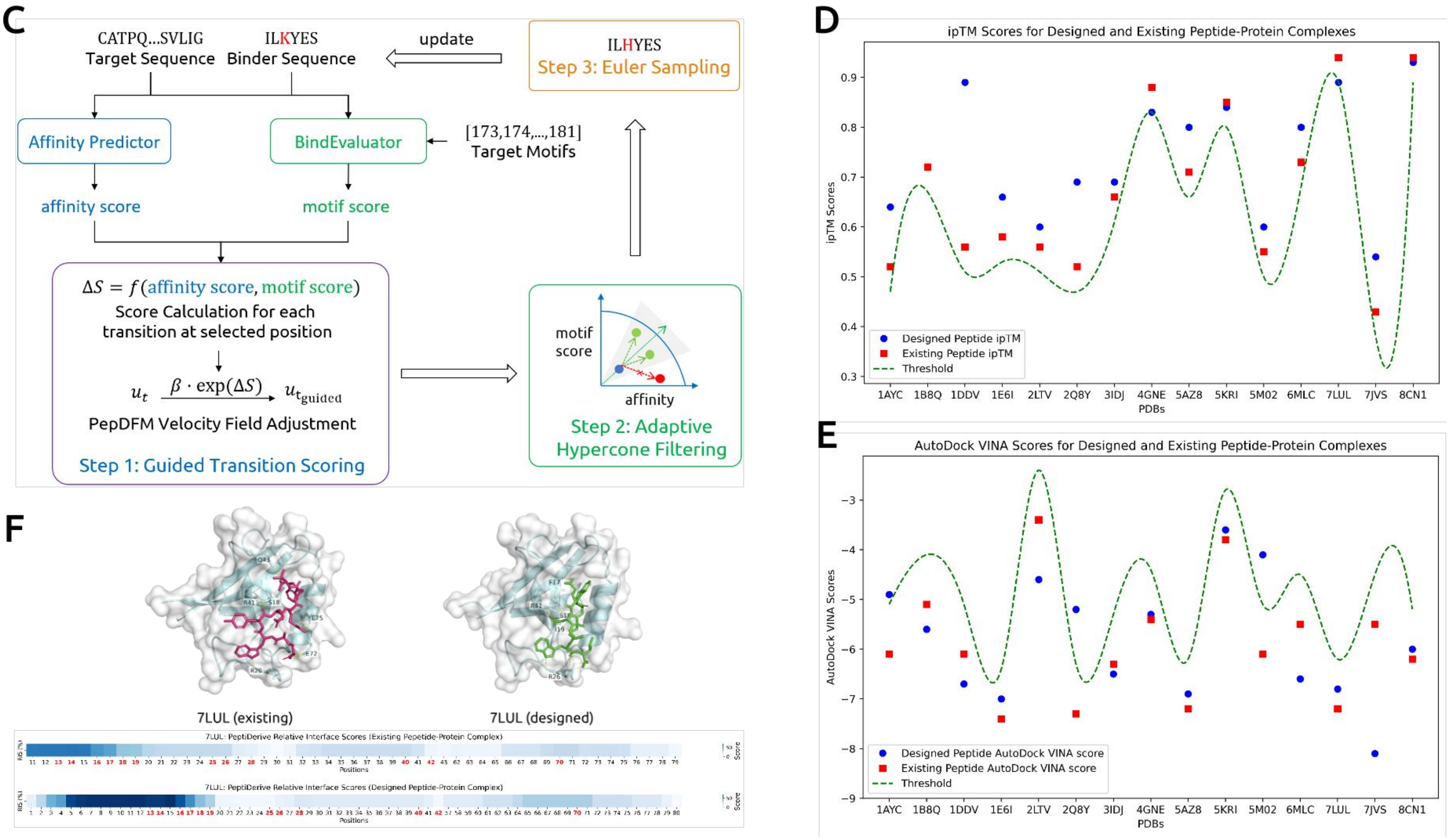
