## Supplementary figures and images for "moPPIt: *De Novo* Generation of Motif-Specific and Functionally Active Peptide Binders via Discrete Flow Matching"

### Figure 2

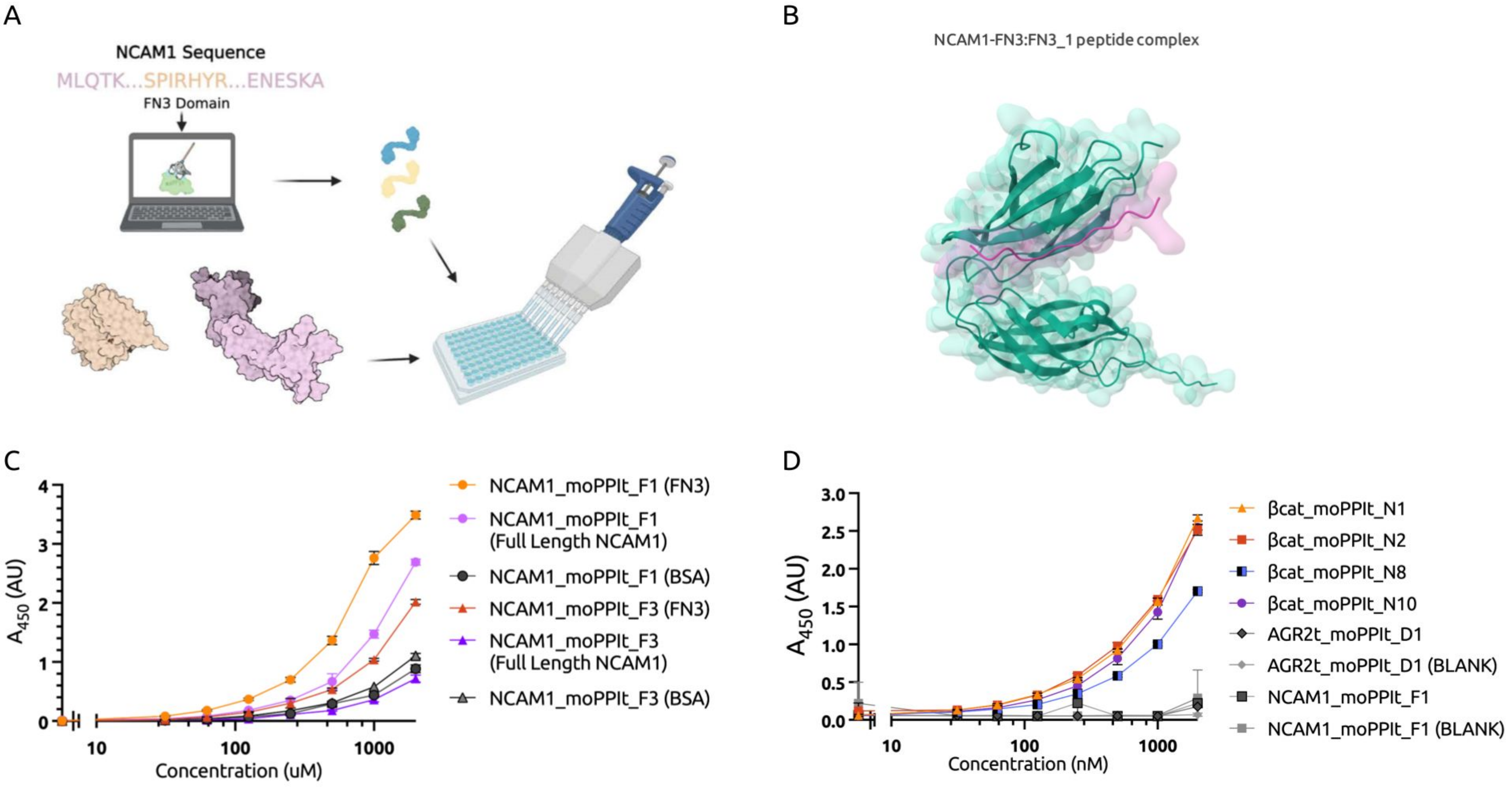

### Figure 3

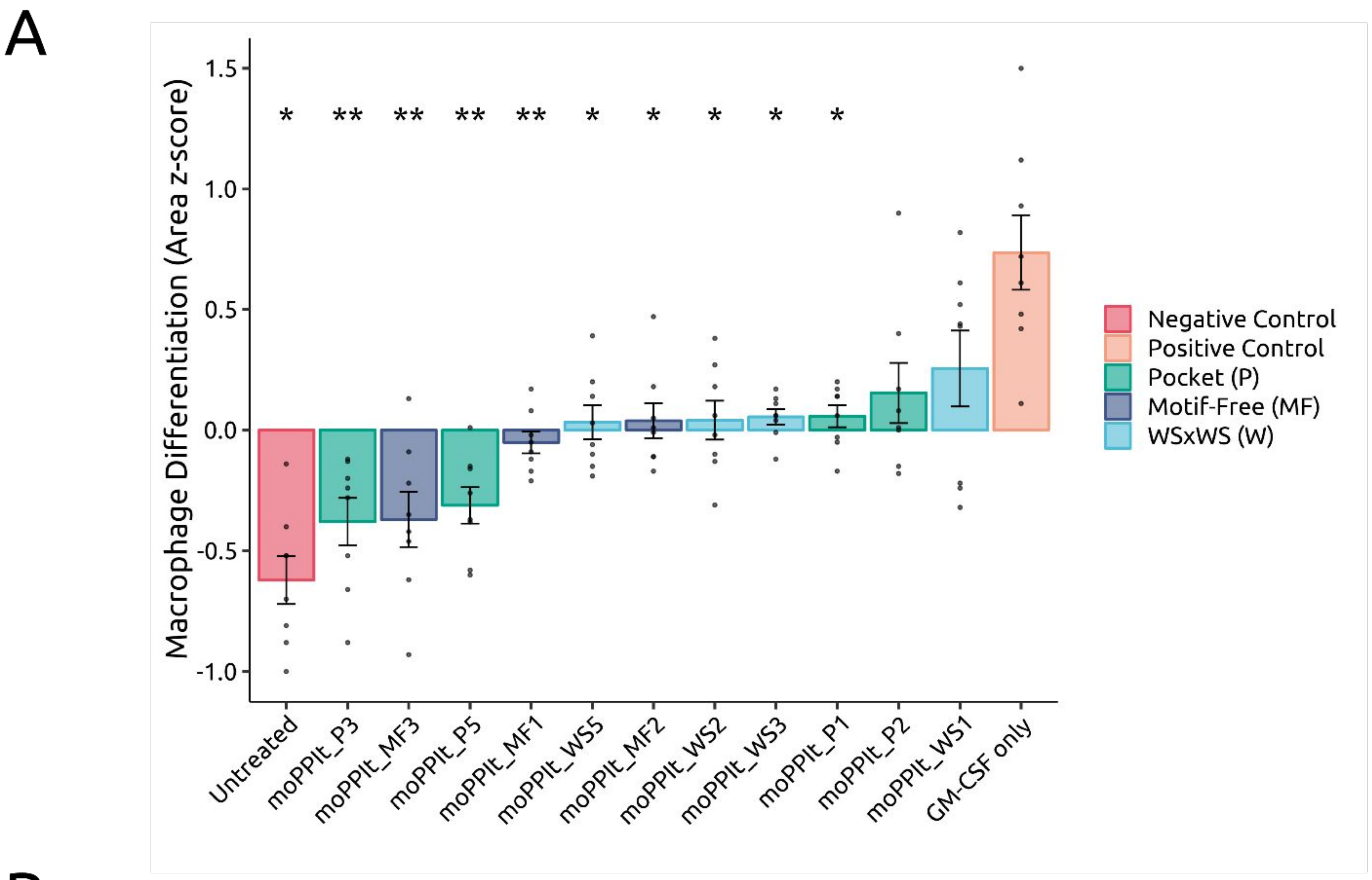

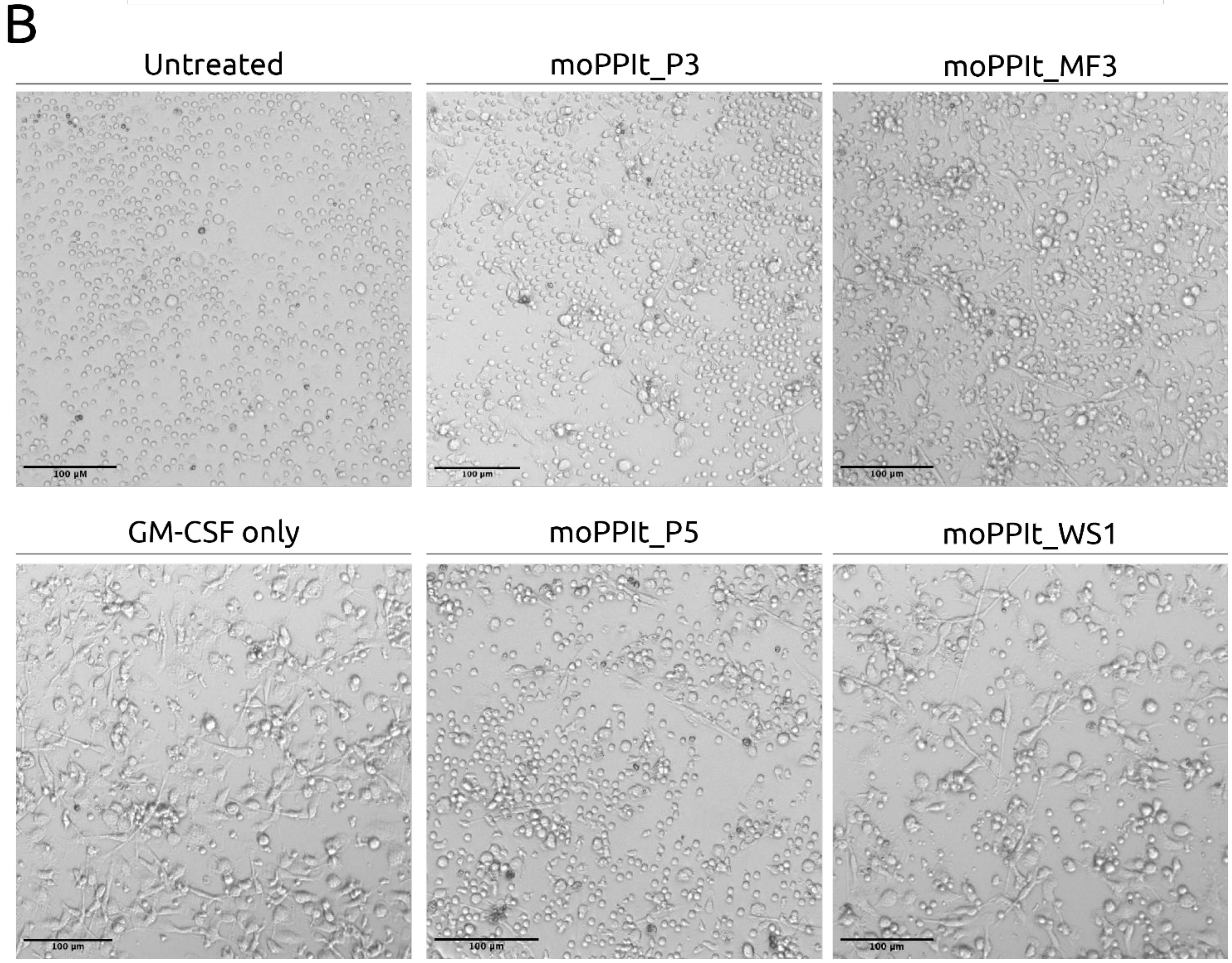

### Figure 4

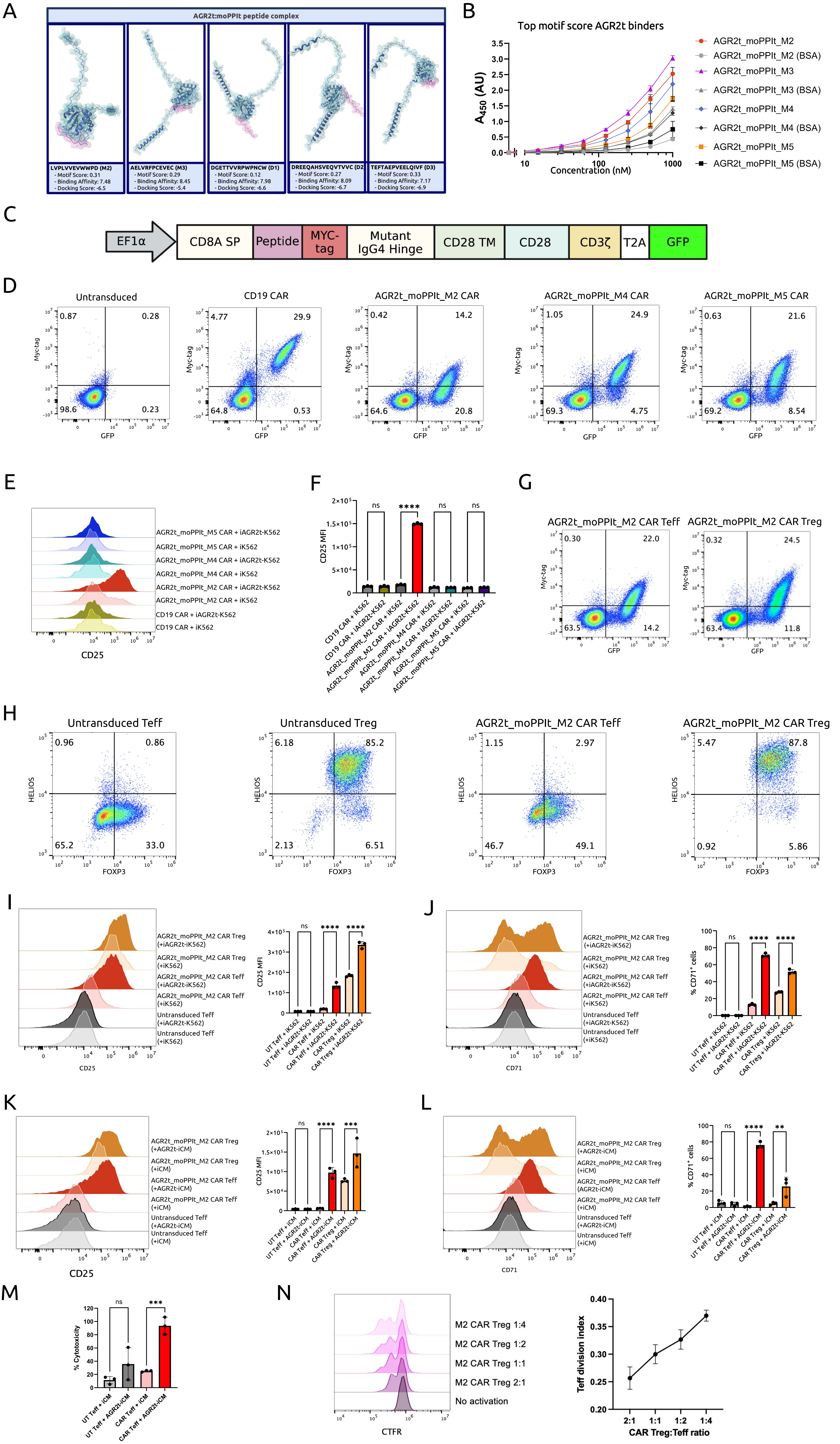
